# RVE based Finite Element Modelling of the Contact Mechanics Between Skin and Indenter

**DOI:** 10.1101/2023.10.01.560340

**Authors:** Rushabh Pardeshi, Naga Neehar Dingari, Beena Rai

**Affiliations:** TCS Research, Tata Research Development and Design Centre, Hadapsar Industrial Estate, Pune 411013, India; TCS Research, Deccan Park, Hyderabad, 500081, India

## Abstract

The nature of contact between human skin and external object (such as medical device, personal care device, fabric and so on) significantly influences the tactile perception and/or the functionality of the object. The contact mechanics depends strongly on the indenter properties and the mechanical properties of skin. Further, the topmost layer i.e., stratum corneum plays the most important role in tactile perception. In this study we use a representative volume element (RVE) based FEM model including the four layers of skin - stratum corneum, epidermis, dermis, and hypodermis - to simulate the contact mechanics between skin and a spherical indenter. The RVE model captures the mechanical properties of the microscopic constituents of the stratum corneum, which is the topmost layer of skin. We found that the RVE model can be used to simulate a variety of stratum corneum conditions (for example, dry and wet stratum corneum) and compositions. Using the RVE model in conjunction with an FEM model, we compute the frictional stress between skin and an indenter, as a function of stratum corneum microstructure, indentation depth, and local coefficient of friction. Both, the RVE model and the contact mechanics model predictions show good qualitative agreement with experimental findings in literature. Such studies will be very useful in in-silico design and optimization of devices that interact with skin. The current framework gives control over several parameters like skin microstructure, indenter or skin geometry and hence can be used to augment/substitute experimental testing.

## 1. Introduction

The skin is the largest organ of the body, and it acts as a boundary between the environment and the internal organs. It is the first line of defense against various mechanical, chemical, thermal or radiation stimuli that may be present in the environment. The skin often comes into contact with various objects such as cloth, footwear, grooming devices and so on. This leads to the development of mechanical stresses on the skin surface because of friction between the skin and object. This may result in discomfort or in worst case, cause skin damage (Gerhardt, Strässle, Lenz, Spencer, & Derler, 2008). Hence, understanding the skin-object contact-mechanics, is important while designing devices/objects that are in contact with the skin.

The stratum corneum (SC), being the topmost layer of skin, has a major role in the contact mechanics between skin and object. Though it has the least thickness (≈20 μ*m*) (Diosa, Moreno, Chica, Villarraga, & Tepole, 2021), it has a considerably higher elastic modulus compared to other layers of skin (Van Kuilenburg, Masen, & Van Der Heide, 2013; Diosa, Moreno, Chica, Villarraga, & Tepole, 2021). Further, its mechanical properties vary over a large range and humidity or water content plays a big role in determining its behavior (Kim & Yun, 2019; Dzidek, Bochereau, Johnson, Hayward, & Adams, 2017; Mojumdar, Pham, Topgaard, & Sparr, 2017; Bernard, et al., 2001; Derler, Rossi, & Rotaru, 2015; Tomlinson, Lewis, Liu, Texier, & Carré, 2011). Increasing the water content has a plasticizing effect on stratum corneum thus making it softer. On the other hand, absence of water from the stratum corneum causes makes it stiffer by orders of magnitude. It is important to consider these effects while characterizing the contact-mechanics between skin and external object. Other skin layers i.e., dermis and hypodermis don’t contribute significantly to the contact mechanics at small indentation depths (∼0-2 mm).

Contact forces are typically composed of two components: normal force, typically described using contact pressure, and the tangential frictional force. It has been found that the Coulomb’s laws (or Amonton’s laws) of friction are not applicable to skin-object interface (van Kuilenburg, Masen, & van der Heide, 2015) to characterize the tangential frictional force. This is because skin is a hyperelastic material (Adams, Briscoe, & Johnson, 2007) whose deformation also contributes to the friction between the skin and external object. The frictional force developed when an object is sliding over the skin has two components – adhesion component and deformation component (Adams, Briscoe, & Johnson, 2007; Pailler-Mattei, Guerret-Piécourt, Zahouani, & Nicoli, 2011; van Kuilenburg, Masen, & van der Heide, 2015). The adhesive component denotes the force required to break the adhesive bonding between the two surfaces. This component depends on the surface properties like interfacial energy, roughness etc. and can be quantified by the product of the shear strength at the interface and the contact area. The second component of the frictional force is due to the elastic deformation that happens during sliding and depends on indentation depth of the object into the skin (Greenwood & Tabor, 1958). In previous works, people have studied the contact properties of skin experimentally, with the help of a spherical indenter. The Hertzian contact theory is a good tool to understand such a contact (Pailler-Mattei & Zahouani, Analysis of adhesive behaviour of human skin in vivo by an indentation test, 2006; Johnson, Kendall, & Roberts, 1971). However, this theory applies only to elastic materials that are homogeneous and isotropic and does not apply to skin, which is a multi-layer material with property variation across the layers.

As the skin-object contact mechanics depends on the mechanical properties of the skin, it varies on account of the inherent variation in the skin properties with age, gender, race, and even anatomical location on the body (Van Kuilenburg, Masen, & Van Der Heide, 2013). Further, extrinsic parameters such as the material properties of the object, temperature and humidity of the environment, and the relative motion between the skin and object also influence the skin-object contact mechanics. Hence, it is difficult to develop a comprehensive analytical model that describes this contact mechanics, by taking into account all the mentioned parameters. Researchers have developed a few empirical models (Veijgen, van der Heide, & Masen, 2013) that consider a few of the control parameters through regression. The regression model usually involves generation of a lot of data by performing in-vivo or in-vitro experiments on human or animal test subject (Pailler-Mattei, Guerret-Piécourt, Zahouani, & Nicoli, 2011; Pailler-Mattei & Zahouani, Analysis of adhesive behaviour of human skin in vivo by an indentation test, 2006; Kim & Yun, 2019; Geerligs, et al., 2011). Over the last few years, researchers worldwide have been moving towards a reduction in animal testing. In certain geographies, there is a ban on the use of animal testing for cosmetic industry applications. Thus, robust in-silico models that can predict the mechanical response of skin are being considered as potential substitutes for animal testing.

Among the various in-silico modelling approaches, finite element models have been extensively used over the past decade to study the biomechanics of skin (Flynn & McCormack, Simulating the wrinkling and aging of skin with a multi-layer finite element model, 2010; Leyva-Mendivil, Page, Bressloff, & Limbert, 2015; Leyva-Mendivil, Lengiewicz, Page, Bressloff, & Limbert, 2017; Santoprete & Querleux, 2014; Nandamuri, 2016; Diosa, Moreno, Chica, Villarraga, & Tepole, 2021) for a variety of applications including wrinkling, subcutaneous drug delivery, wound healing and robotics assisted surgery. Recent advancements in the development of commercial and open-source FEM packages have allowed researchers to develop robust in-silico models of skin that can a) include property variation across different skin layers b) model contact mechanics between skin and indenter for large deformations and include microscopic details of skin structure into the modelling framework. Previous works (Leyva-Mendivil, Lengiewicz, Page, Bressloff, & Limbert, 2017; Leyva-Mendivil, Page, Bressloff, & Limbert, 2015) have characterized the influence of skin topography on skin mechanics using FEM framework. Further, using Representative Volume Elements (RVE) (Santoprete & Querleux, 2014; Nandamuri, 2016) one can include even the microscopic (lower length scale) properties into the macroscopic finite element modelling framework. While RVE based models are extensively used in the context of hard composites, their application to soft materials like skin (Santoprete & Querleux, 2014) has been explored only in the recent past. Owing to the multiscale structure of the skin, RVE approach is a very efficient way to incorporate the complex lower-length scale effects to simulate the macroscopic mechanical response.

In this study, we present an RVE based FEM model to characterize the contact mechanics between the skin and a rigid spherical indenter. The variation in stratum corneum properties which play the most important role in the contact mechanics is modelled using the Representative Volume Element (RVE) methodology, which allows us to include the properties of micro-constituents such as corneocytes, corneodesmosomes and lipid matrix into the modelling framework. Further the variations in skin properties across different layers can be easily considered using the multi-layer geometrical representation in the FEM framework. For example, the skin thickness may vary with anatomical location, race or age of the subject and this can be easily changed in the FEM model. Similarly, the material properties or the geometry of the indenter can also be varied. This provides a framework where multiple permutations and combinations can be applied to the skin and indenter properties, as well as to the relative motion between the two. A lot of data can thus be generated in-silico, which can aid in the design of any device or other objects which are constantly in contact with skin and improve its functionality or comfort level.

In section 2 we discuss the FE framework and the RVE approach used to develop the multilayer skin model and the contact simulations. The results are discussed further in section 3.

## 2. Mathematical modelling

In this section, we describe the representative volume element (RVE) based finite element (FE) framework that is used to study the contact mechanics between skin and indenter. Broadly speaking, skin has three different layers – the epidermis, dermis and hypodermis. The epidermis can be further divided into two regions: the stratum corneum and viable epidermis. The human skin is generally modelled as a multi-layer material with different mechanical properties for different layers (Flynn & McCormack, Simulating the wrinkling and aging of skin with a multi-layer finite element model, 2010; Leyva-Mendivil, Page, Bressloff, & Limbert, 2015) because of considerable variation in the properties across different layers. As explained in the previous section, stratum corneum plays a significant role in the contact mechanics between skin and indenter despite its small thickness because of its high elastic modulus compared to the other skin layers (Leyva-Mendivil, Page, Bressloff, & Limbert, 2015). Further, the mechanical properties of stratum corneum strongly depend on the properties of its micro-constituents and hydration state (Santoprete & Querleux, 2014). Thus, the mechanical properties of stratum corneum are described using RVE based approach (section 2.1.1), while those of inner skin layers are described using hyperelastic constitutive models.

### 2.1 Constitutive modelling of stratum corneum

#### 2.1.1 Structure of the stratum corneum

As shown in figure 2, stratum corneum consists of alternating layers (around 10-15) of two planar (very thin compared to other dimensions) materials. The first planar material is composed of planar hexagonal corneocytes (brick) separated by peripheral intercellular space (mortar). The second planar material is composed of non-peripheral intercellular space (mortar). Both peripheral and non-peripheral intercellular spaces consist of corneodesmosome fibers embedded in a lipid matrix. Thus, the stratum corneum can be geometrically represented by two RVEs: a) cellular scale or first RVE and b) macroscale RVE or second RVE.

**Figure 1:**
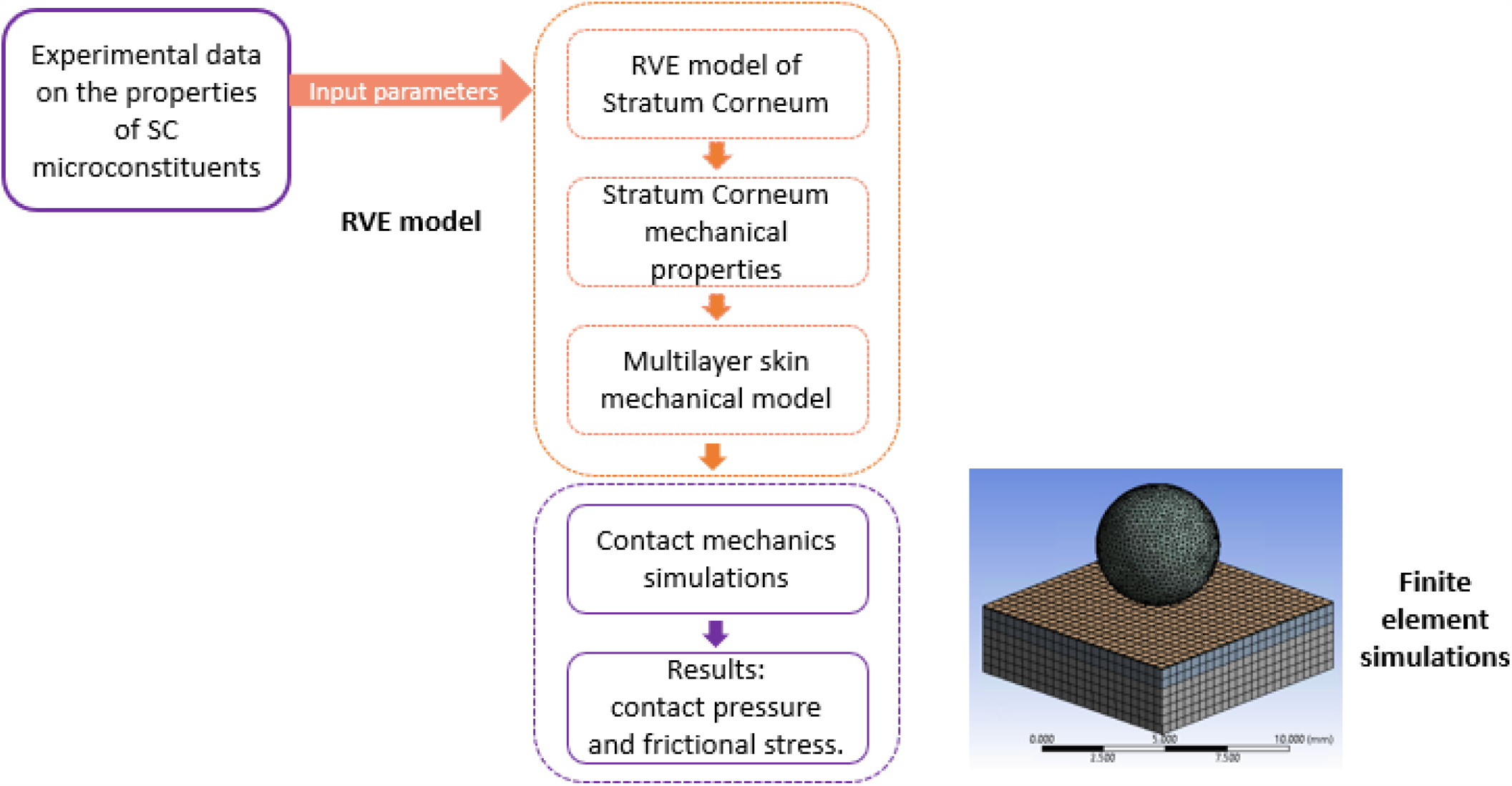
An RVE based FEM framework for modelling the contact mechanics between skin and a device. The framework shows how the properties of the microconstituents of stratum corneum can be represented in the finite element modelling of contact mechanics using an RVE based approach.

**Figure 2:**
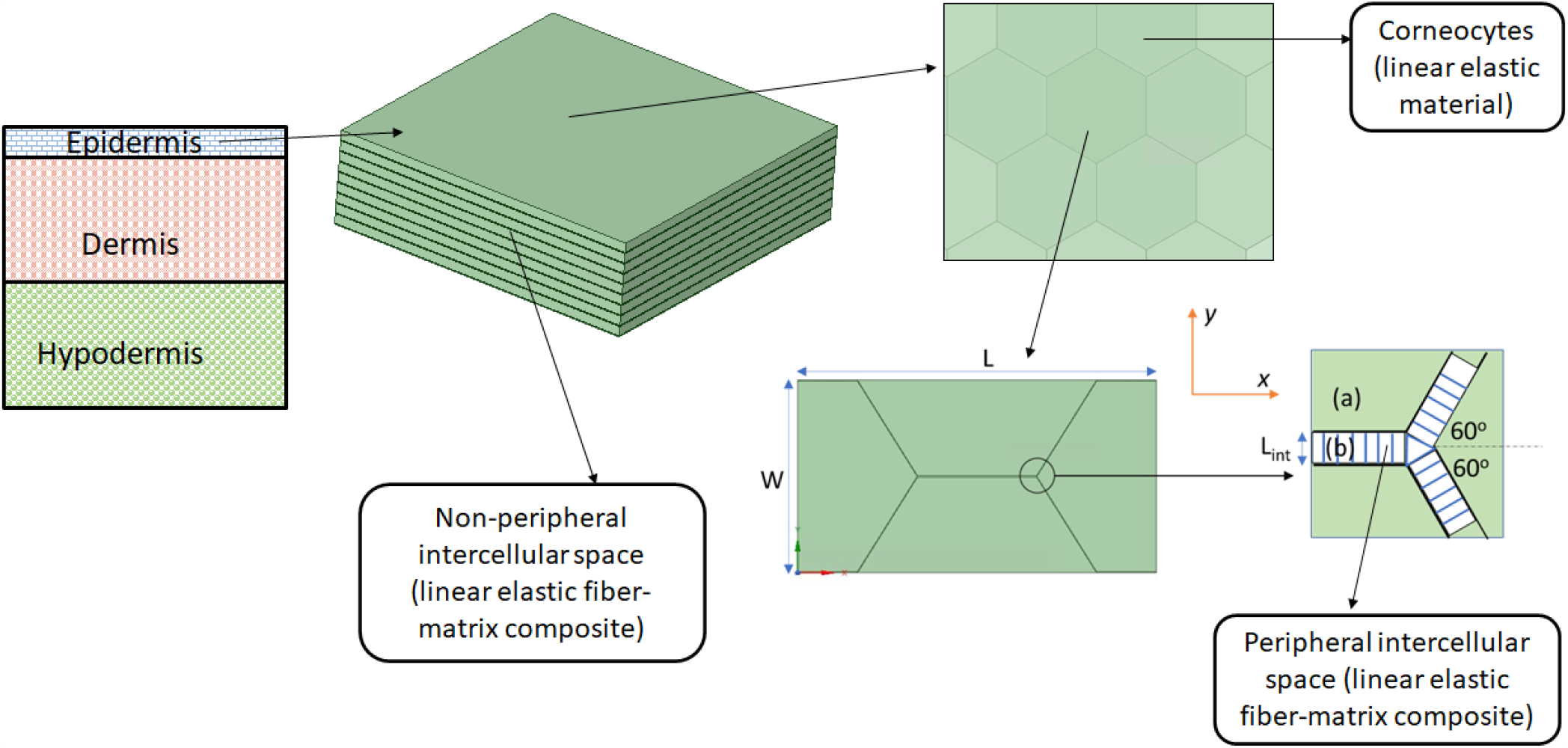
The hexagonal structure of the Stratum Corneum (SC) at cellular scale. It is like a brick-and-mortar structure, with the corneocytes being the main structural component and lipid-corneodesmosome composite acting like mortar.

The geometry of the first RVE/cellular RVE is shown in figure 3. The hexagonal corneocytes are much larger in width compared to the (peripheral) intercellular space surrounding them. The intercellular space is a lipid-corneodesmosome composite material, with corneodesmosome fibers embedded in the lipid matrix. The corneodesmosome fibers oriented perpendicular to the corneocyte surfaces (figure 3, zoomed in view). The volume fraction of corneodesmosome within the intercellular space typically varies depending on the skin composition.

**Figure 3:**
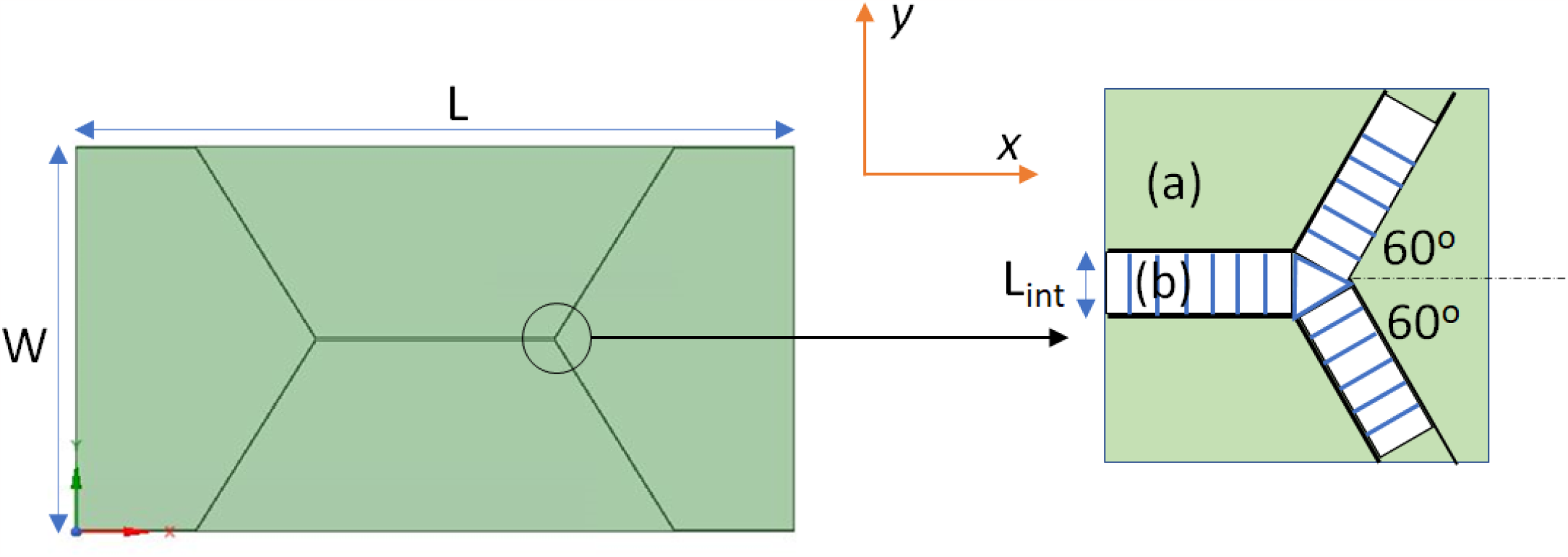
The first RVE: The cellular scale structure of the stratum corneum can be represented by an RVE as shown here. L = 60 µm, W = 34.64 µm, L_int_ = 0.03 µm. The corneocytes (a) take up the most space. The space between corneocytes (b) is occupied by a composite of lipids and corneodesmosomes. It can be represented as a fiber (corneodesmosomes) – matrix (lipids) composite, corneodesmosomes always being perpendicular to the corneocyte edges as shown in the figure.

To obtain the entire stratum corneum structure, one needs to stack up multiple layers of the first RVE, as shown in figure 4. The space between these layers (non-peripheral) is filled up by the lipid-corneodesmosome composite, with the fibers oriented perpendicularly to the layers. The cellular layer thickness is typically ≈0.3 µm and the intercellular space thickness is ≈0.03 µm.

**Figure 4:**
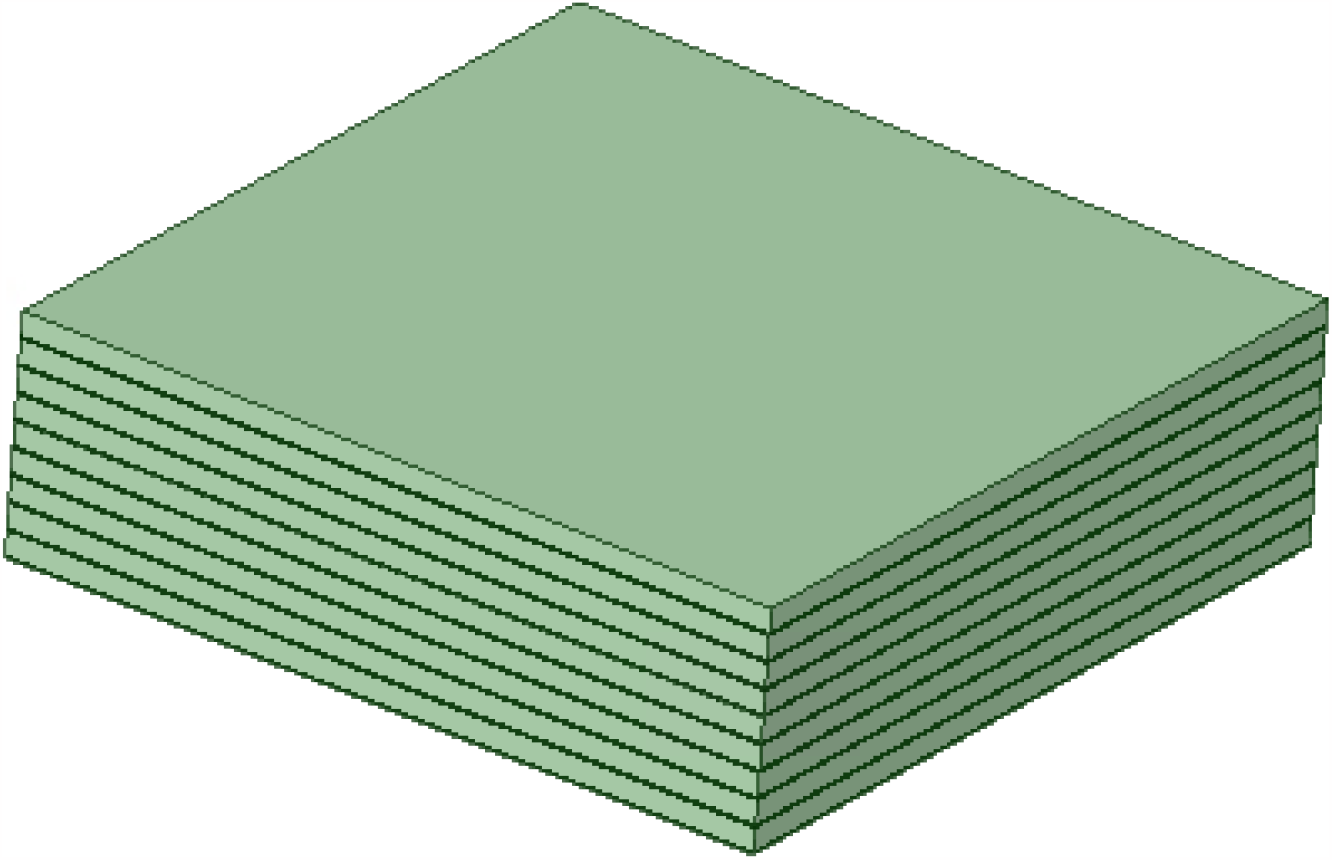
The second RVE: The cellular scale layers from the first RVE are stacked over each other to form the second RVE. The space between these layers is also occupied by the lipid-corneodesmosome composite, with the fibers perpendicular to the cellular layers. This structure is similar to the stratum corneum structure, which is made up of 10-15 cellular layers.

#### 2.1.2 Computation of stratum corneum elasticity

We assume that the global coordinate system xyz is aligned such that the first RVE is in the xy plane and the z-axis is normal to the planar first RVE. Corneocytes are assumed as isotropic and linear elastic (Elasticity *E*_*c*_, Shear modulus *G*_*c*_ and Poisson’s ratio *vc*); and the intercellular spaces are assumed to be described as fiber-matrix composites. Since the corneodesmosome fibers in the intercellular spaces are oriented perpendicular to the surface of corneocytes, their orientation with respect to global coordinate system changes with position. Thus, the computation of elasticity tensor of intercellular space is non-trivial. In case of non-peripheral intercellular space, the fibers are oriented along the z-axis. Thus the computation of elasticity tensor for non-peripheral RVEs is relatively simpler and can be obtained using mixture theory and orthotropy assumption (Santoprete & Querleux, 2014).

To facilitate the computation of elasticity tensor for peripheral intercellular space, we define a local coordinate system (1-2-3) with direction 1 oriented along the direction of fibers and directions 2,3 normal to it (Figure 5). The elasticity tensor in the local coordinate system can be obtained by mixture theory (Santoprete & Querleux, 2014). This elasticity tensor in the local coordinate system needs to be mapped to the global coordinate system xyz using coordinate transformation procedure as outlined by (Daniel, Ishai, Daniel, & Daniel, 2006).

**Figure 5:**
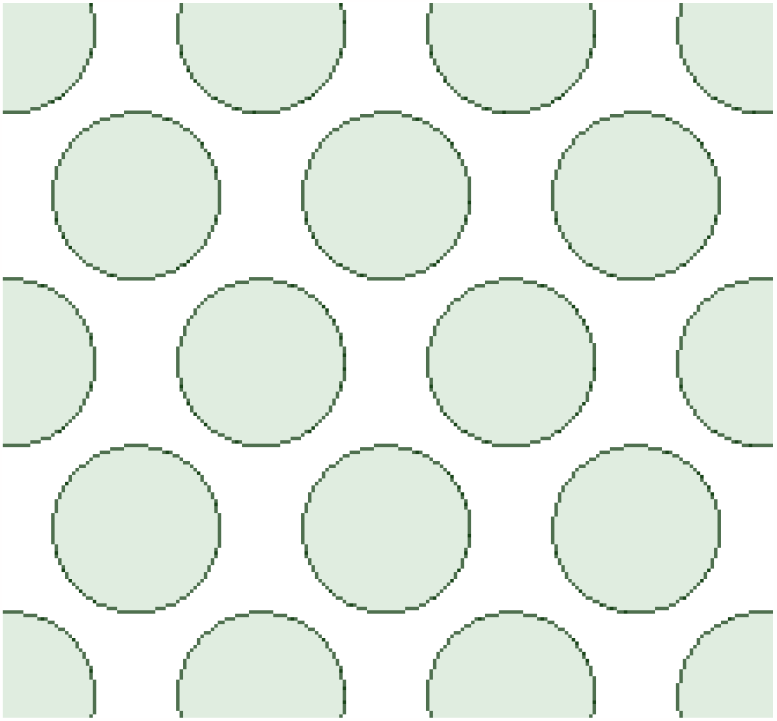
The plane of isotropy (2-3) in a fiber-matrix composite RVE, with the fibers oriented in direction 1. Here, lipid is the matrix and corneodesmosomes are the fibers.

##### Elasticity tensor in the local coordinate system

In the local coordinate system 1-2-3, the components of the elasticity tensor are given as (Santoprete & Querleux, 2014)

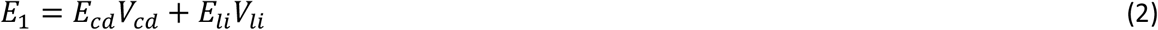

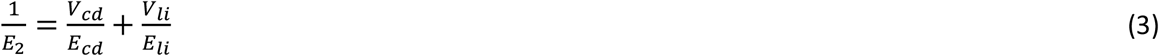

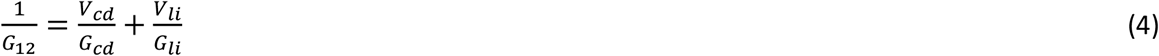

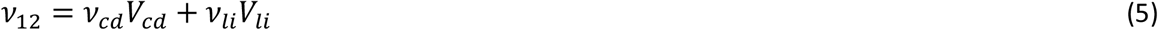

By assuming symmetry in the directions 2,3, we have *E*_3_=*E*_2_,*G*_13_=*G*_12_,*V*_13_=*V* _12_. Here E_cd_ and *Eli* are the elasticities of corneodesmosomes and lipid matrix respectively; *G*_*c*d_ and *G*_*li*_ are the corresponding shear moduli; and *v*_*c*d_ and *vli*are the Poisson ratios. *G*_23_and *v*_23_cannot be directly obtained from the mixture theory In order to find *G*_23_and *v*_23_, a simple fiber-matrix RVE with fibers oriented in direction 1, and with lipid as matrix and corneodesmosomes as fiber (with an assumed volume fraction) was generated in ANSYS and solved for resultant elasticity parameters by applying periodic boundary condition (described later). From the output of this RVE, we can find *G*_23_and *v*_23_, which are the shear modulus and Poisson’s ratio for the plane of isotropy. The compliance components in the local coordinate system are given in terms of the elasticity tensor components as (Daniel, Ishai, Daniel, & Daniel, 2006)

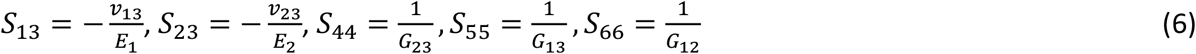

##### Elasticity tensor in the global coordinate system

The in-plane components of the elasticity tensor in the global coordinate system xyz are given as

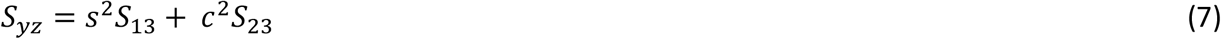

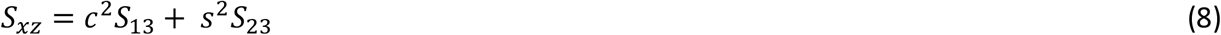

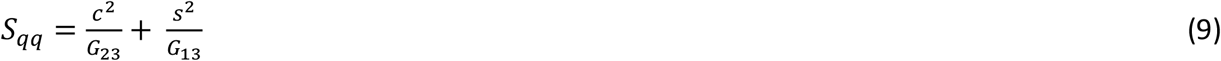

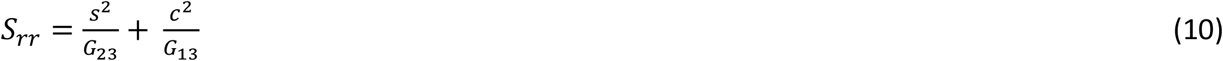

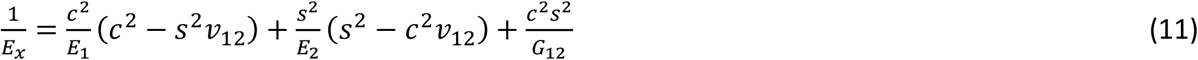

Here, ‘s’ and ‘c’ represent sine and cosine of fiber orientation θwith respect to the X-axis in the XY plane (Figure 6). Further, the out of plane the shear modulus and Poisson ratio in the global coordinate system are given as

**Figure 6:**
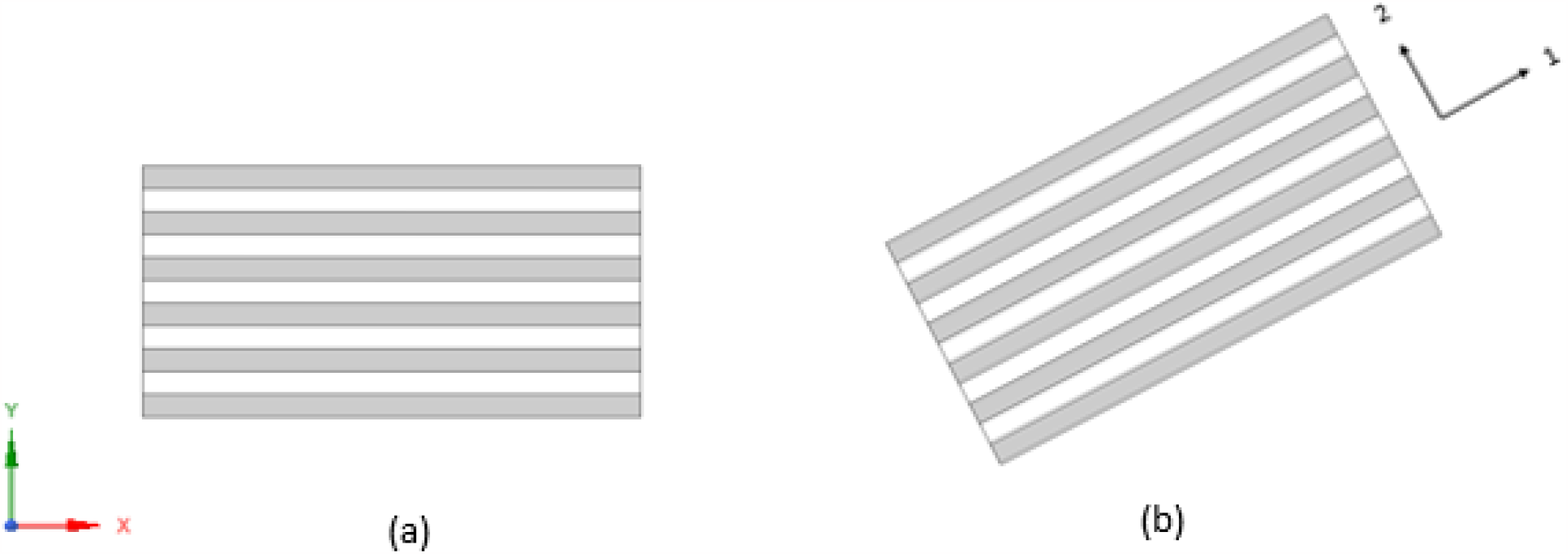
Using mixture theory, the equivalent elasticity of a fiber matrix composite in terms of its constituents can be calculated if the fiber orientation is in global X-Y orientation. However, if fibers are oriented locally in a different 1-2 direction, at some angle with respect to global X-Y direction, we need to apply transformation equations to calculate equivalent elasticity in X-Y direction.

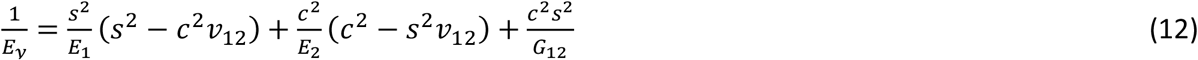

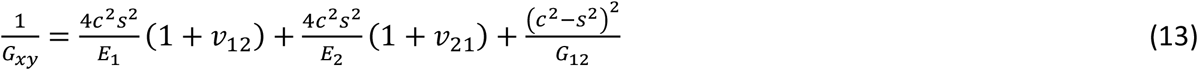

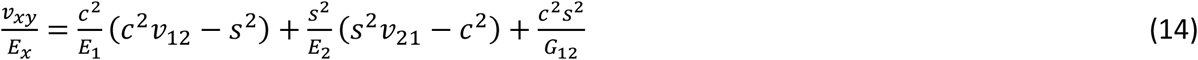

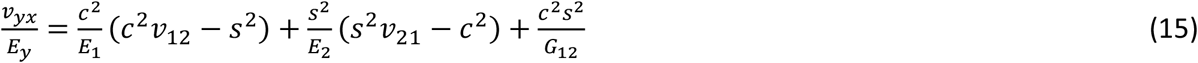

Where the compliance tensor components *S*_*xz*_, *S*_*yz*_, *S*_*qq*_, *S*_*rr*_ are given as

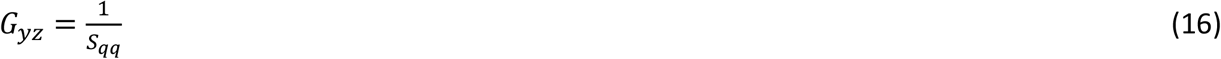

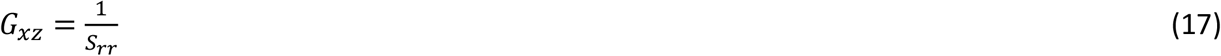

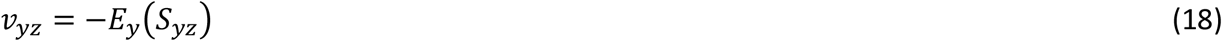

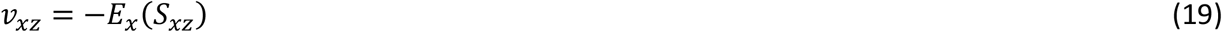

#### 2.1.3 Homogenization to obtain the effective properties of stratum corneum

The overall elasticity of stratum corneum is obtained from the elasticity tensor of individual components using RVE homogenization technique. This can be done by applying one of the following boundary conditions to the RVE: uniform displacement, uniform traction, or periodic boundary condition. For periodic structures (like stratum corneum RVE), using the periodic boundary conditions (PBC) satisfies the displacement as well as traction continuities. Also, the elasticities computed using PBC show no dependence on the window size (number of periodic unit cells considered) of the RVE (Nandamuri, 2016).The constraint equation applied at two opposite boundaries can be represented in terms of displacement differences as (Nandamuri, 2016):

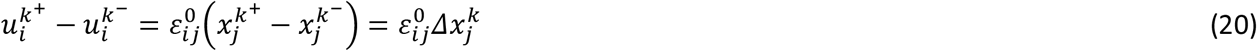

Here, 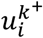 anmacroscopic uniform strain on the boundaries. More d 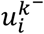 are the displacements at k^th^ pair of opposite boundaries and 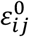 is the applied macroscopic uniform strain on the boundaries. More details on PBC and its application in finding homogenized properties of an RVE can be found in (Nandamuri, 2016). For the first RVE, we identified periodicity in XY plane (figure 3) and applied periodic boundary conditions to solve for elastic constants. These elastic constants were taken as inputs for the second RVE, and in similar fashion, periodic boundary conditions were applied to the second RVE to get the overall stratum corneum’s elastic properties. This procedure was done with the help of Ansys 19.2.

##### 2.2 Constitutive modelling of other skin layers

The viable epidermis, dermis and hypodermis are assumed to be described by Ogden hyperelastic model with the following strain energy density function:

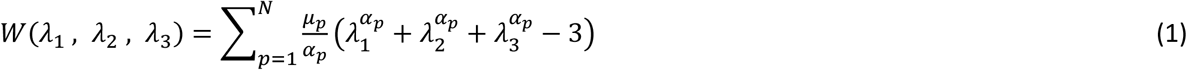

Here, λ_n_ are the stretch ratios and N, α, µ are the material constants. We assumed a first order Ogden model and the material constants are mentioned in the subsequent section.

### 2.2 Contact mechanics

As noted in the introduction section, frictional force can be due to adhesive forces at the interface or due to mechanical deformation. To simplify things, we consider only the influence of mechanical deformation and neglect the adhesion component of friction in this study. The contact mechanics in finite element framework is described by identifying a) the contact type (node to surface/surface to surface) to estimate the extent of contact between two surfaces and b) the contact algorithm to describe the interfacial tangential and normal forces between the two surfaces. For the latter, one can choose either Lagrangean contact algorithm or Penalty algorithm (Ansys Contact Technology Guide, 2009; Weyler, Oliver, Sain, & Cante, 2012) or a combination of these two like augmented Lagrangean contact algorithm. Both these features are readily available in typical finite element solvers like Ansys Workbench.

## 3 Results and Discussion

In this section, we discuss the influence of the properties of microconstituents of the stratum corneum on its elastic properties (sec 3.1). We present two cases in which the overall elasticity of the stratum corneum roughly corresponds to that of dry and wet skin respectively reported in the literature. Next in sec 3.2 we present the results of the contact mechanics simulations using a spherical indenter for the two cases mentioned above.

### 3.1 RVE model of Stratum Corneum

The mechanical properties of the three components, lipids, corneocytes and corneodesmosomes, are taken from the values reported in the literature (Santoprete & Querleux, 2014). The mechanical behavior of all the three components was assumed to be in the linear elastic regime. The overall elasticity of the stratum corneum was found to vary strongly with change in the elasticity of corneodesmosomes and the lipids. Thus we have varied these two properties till the effectivity elasticity (discussed in figure 11) roughly corresponded to that of dry skin (421.99 MPa) and wet skin (9.66 MPa). In literature, the values of stratum corneum elasticity modulus for dry and wet skin are assumed to be 350 MPa and 7 MPa respectively (Leyva-Mendivil, Lengiewicz, Page, Bressloff, & Limbert, 2017). Thus, our values of effective elasticity of the stratum corneum roughly correspond to the dry and wet skin conditions. The mechanical properties of the components in the RVE as shown in table 1 below.

**Table 1:** Properties of the individual components of the RVE, for two different cases of stratum corneum elasticity. E_c_ = corneocyte elasticity, E_cd_ = corneodesmosome elasticity, E_li_ = lipid elasticity, v = Poisson’s ratio, V = volume fractions within the intercellular space. These properties were used for both the RVEs. These are approximate values taken from the literature (Santoprete & Querleux, 2014)

| Property | Dry skin ($E \approx 421.99$ MPa) | Wet skin ($E \approx 9.66$ MPa) |
| --- | --- | --- |
| $E_c$ | 1 GPa | 1 GPa |
| $v_c$ | 0.2 | 0.2 |
| $E_{cd}$ | 2.5 MPa | 30 kPa |
| $E_{li}$ | 0.25 MPa | 3 kPa |
| $v_{cd}$ and $v_{li}$ | 0.2 | 0.2 |
| $V_{cd}$ and $V_{li}$ | 0.5 | 0.5 |

#### Stratum Corneum elasticity using RVE model

The elasticity of intercellular space plays a crucial role in the overall elasticity of the stratum corneum. Intercellular space is a fiber matrix composite which can be either peripheral or non-peripheral as discussed in section 2.1.2. In the peripheral intercellular space, the corneodesmosome fibers can be oriented at either 90, 30 or 150 degrees with respect to x-axis depending on whether the corneocyte edge is oriented at 0, 60 or 120 degrees respectively with the x-axis. Using the methodology described in section 2 (equations 4-22), the elasticity tensor of the intercellular space is evaluated and the results are summarized in figures 7-10 for the microconstituent properties corresponding to the dry and wet stratum corneum. We assume the same global coordinate system as described in section 2.1.2 to discuss the results.

**Figure 7:**
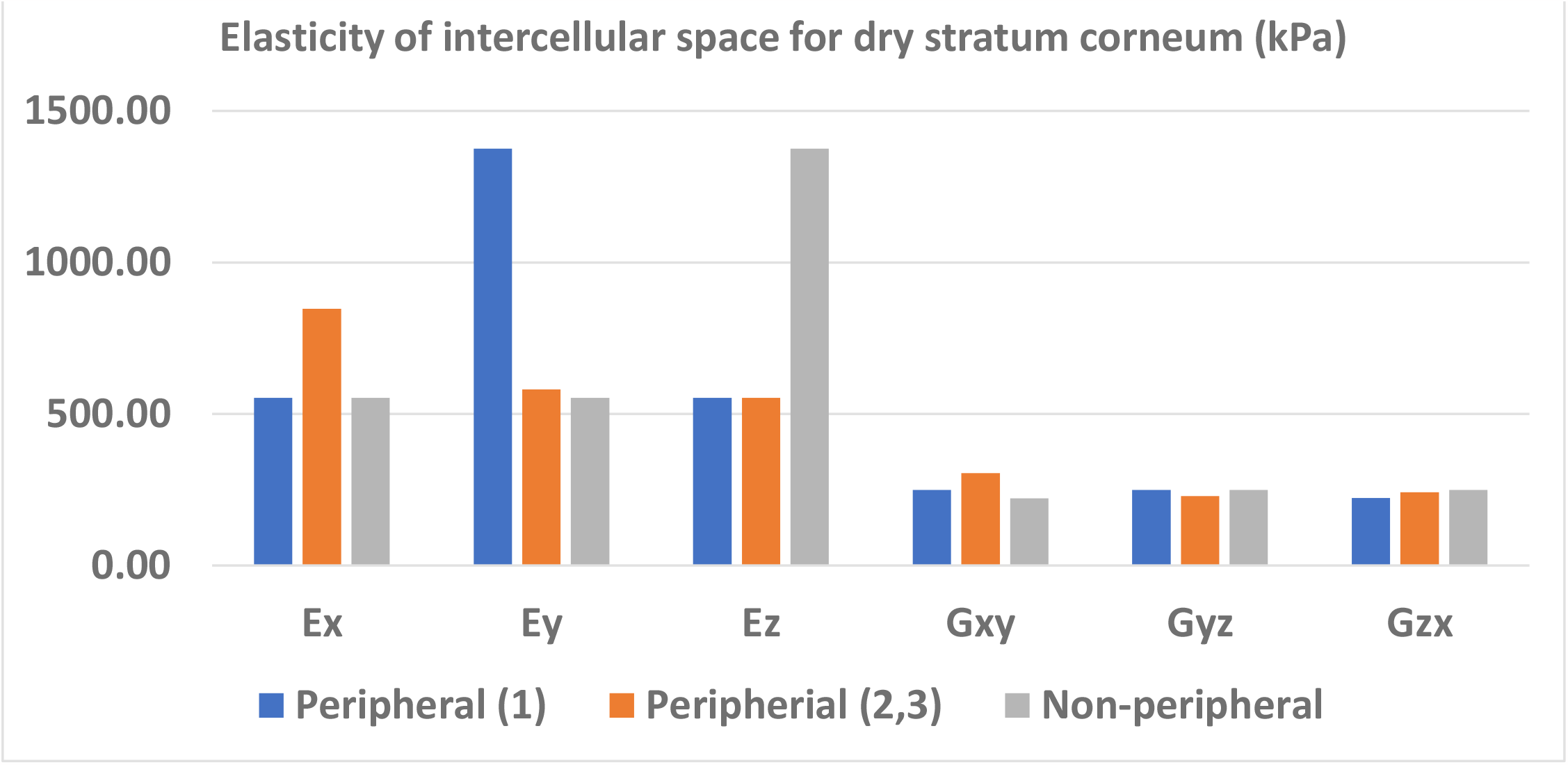
The elastic constants of the peripheral(1,2,3) and non-peripheral intercellular spaces with respect to the global XYZ coordinate system, for dry skin, calculated using equations (2-19).

In what follows, Peripheral (1), Peripheral (2) and Peripheral (3) refer to the intercellular spaces in which fibers are oriented at 90, 30, 150 degrees respectively with respect to the global x-axis ( as shown in figure 3). It can be observed from figures 7 and 9 that when the fibers are oriented in any of the global coordinate directions (Peripheral (1) and Non peripheral), the elasticity in the direction of fiber orientation is higher. This is because in any fiber matrix composite, the material is stronger in fiber direction. The elastic properties for peripheral 2 and peripheral 3 intercellular spaces are similar because of symmetry. It can also be seen in figures 7-10 that for peripheral(1) intercellular space *E*_*x*_=*E*_*z*_, *G*_*xy*_ = *G*_*yz*_, *v*_*xy*_ =*v*_yz_. This is because XZ plane which is normal to the fiber orientation is the plane of isotropy for peripheral(1) intercellular space. Similar observations can be made for non-peripheral intercellular space, where XY is the plane of isotropy. From equation 5, we see that the Poisson’s ratio of a composite depends on the volume fractions and individual Poisson’s ratio of components. Since we kept the volume fraction of lipid and corneodesmosomes at 0.5 each and used similar Poisson’s ratio for both (0.2), the overall Poisson’s ratio for each case in figures 8 and 10 is also close to 0.2. Comparing figures 7 and 9, we see that the overall elasticity values for intercellular space in wet stratum corneum is very less compared to dry stratum corneum. Thus the RVE model has the flexibility to incorporate a variety of skin conditions by the appropriate choice of the microconstituent properties.

**Figure 8:**
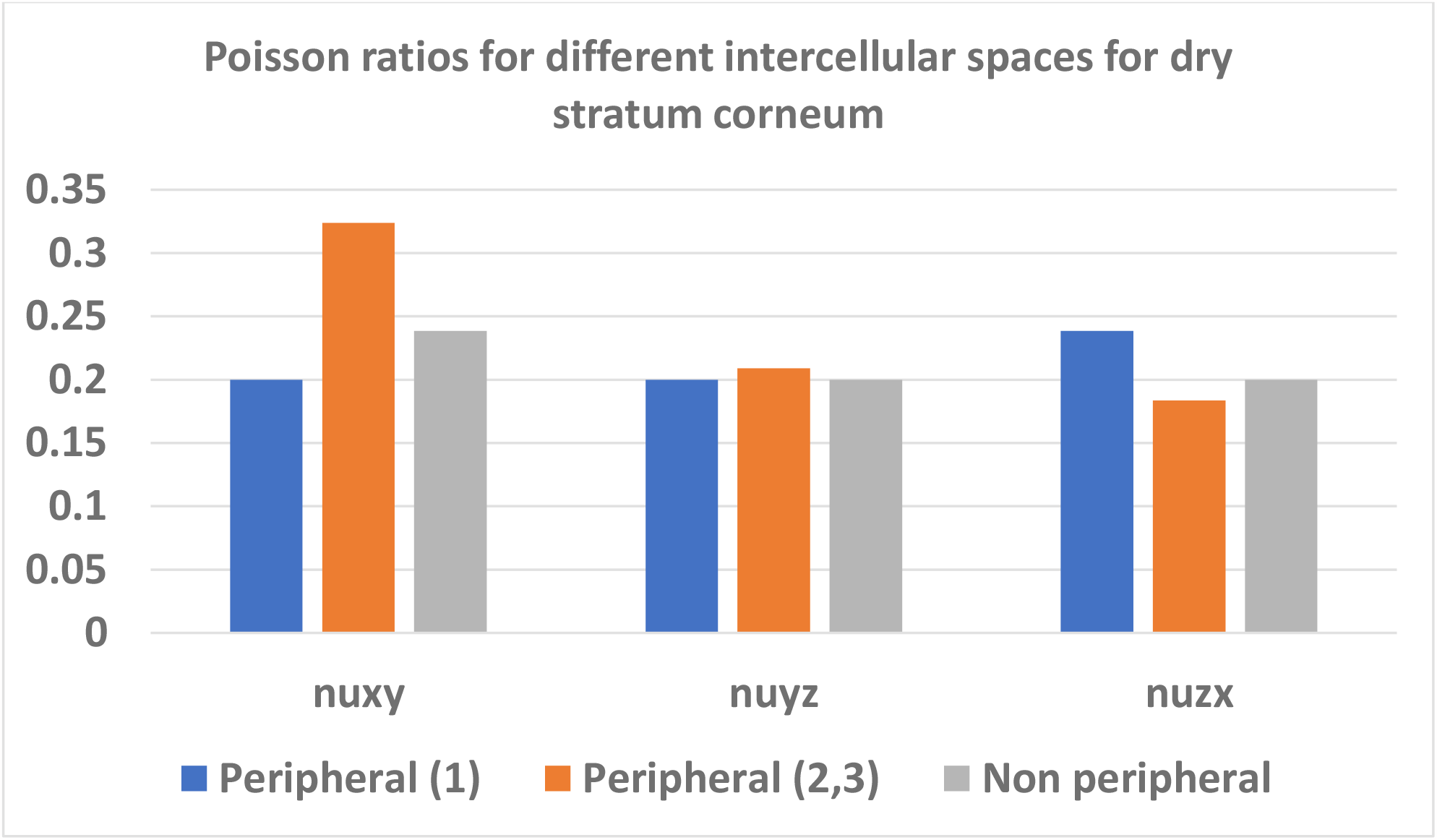
The Poisson ratios of the peripheral(1,2,3) and non-peripheral intercellular spaces with respect to the global XYZ coordinate system, for dry skin, calculated using equations (2-19).

**Figure 9:**
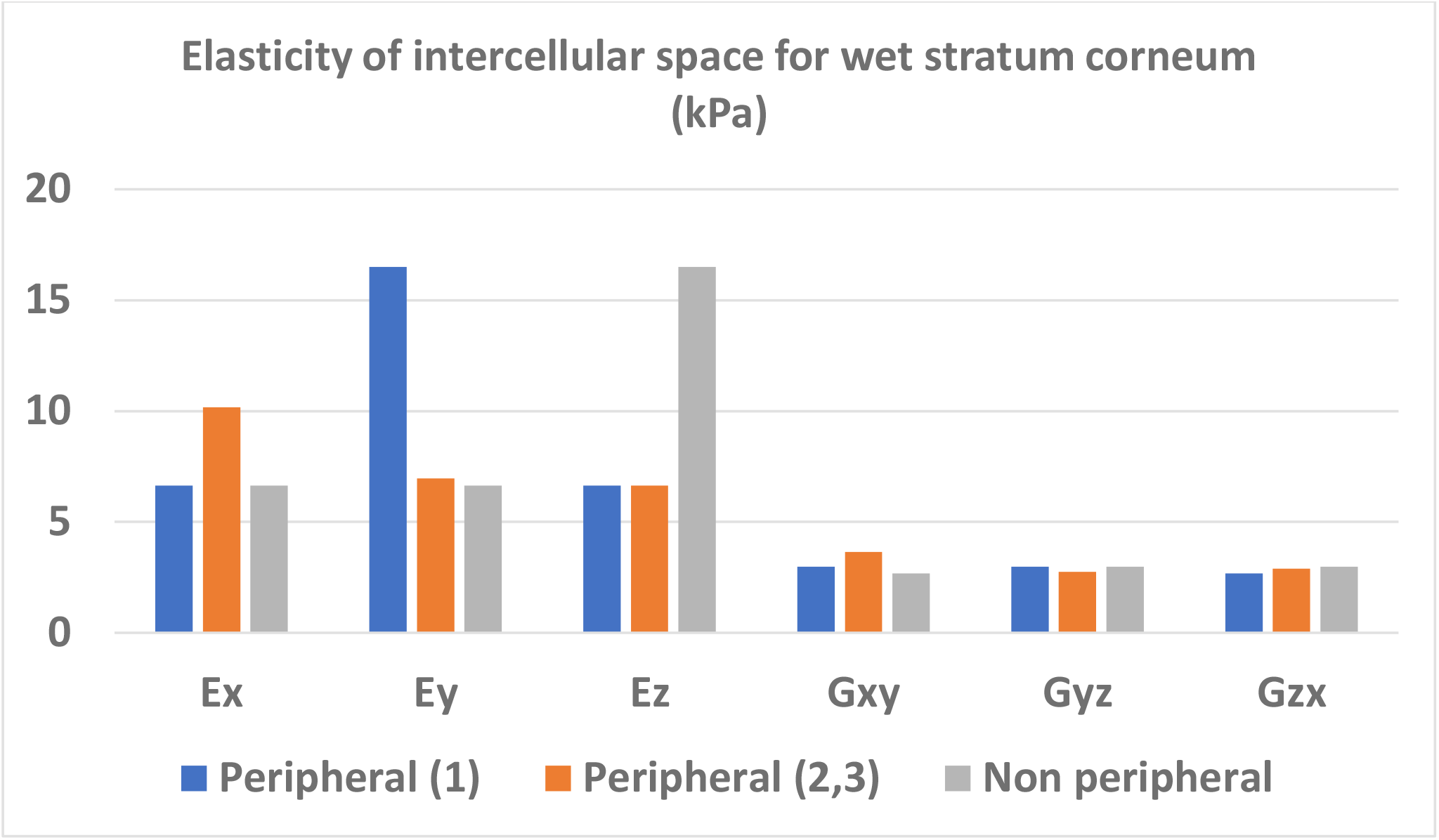
The elastic constants of the peripheral(1,2,3) and non-peripheral intercellular spaces with respect to the global XYZ coordinate system, for wet skin, calculated using equations (2-19).

**Figure 10:**
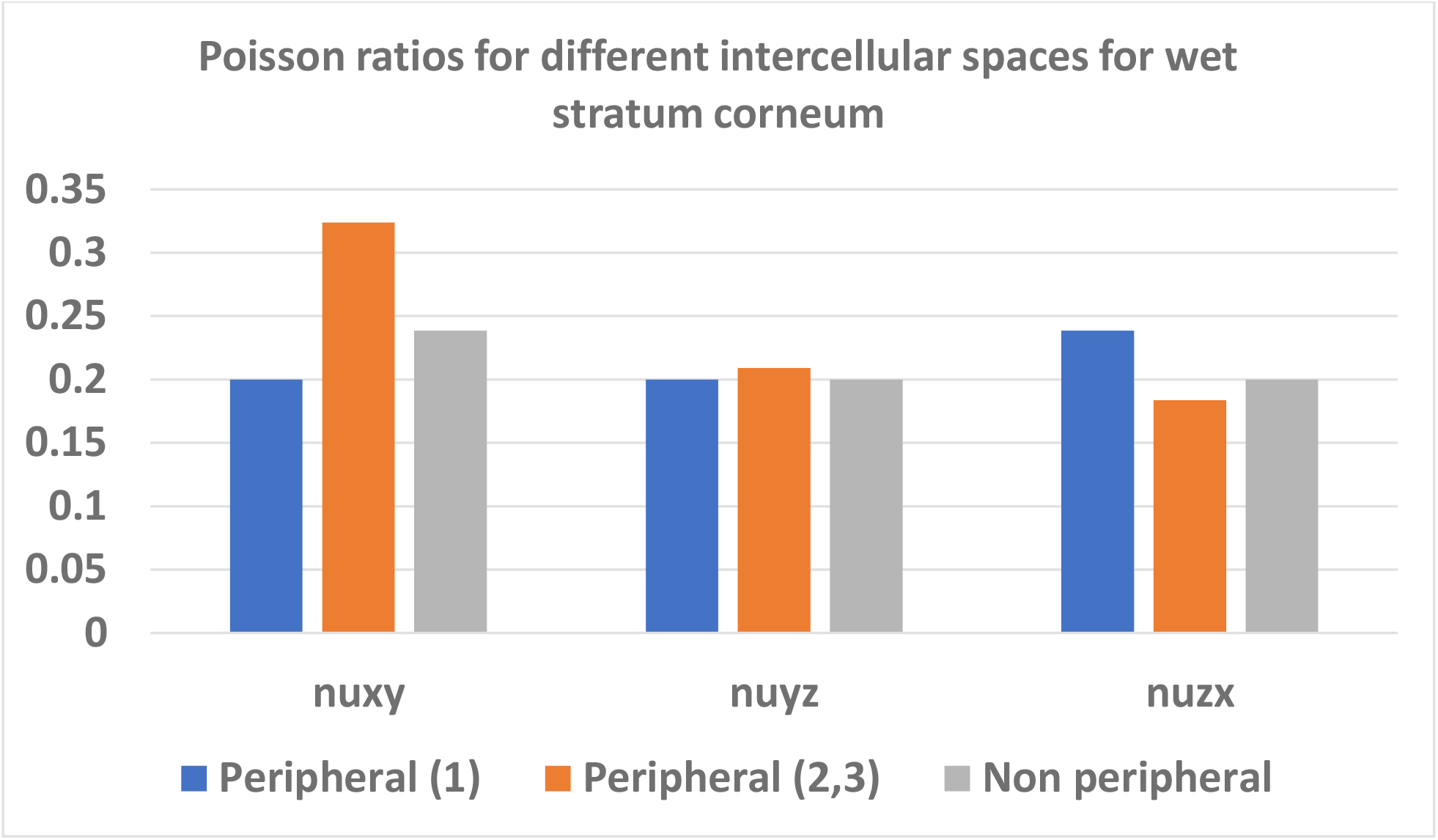
The Poisson ratios of the peripheral(1,2,3) and non-peripheral intercellular spaces with respect to the global XYZ coordinate system, for dry skin, calculated using equations (2-19).

After computing all the required elastic constants of the peripheral and non-peripheral intercellular spaces, the values were given as inputs to both the RVE models to evaluate the overall elasticity of the stratum corneum. Corneocytes are modeled as linear isotropic, and their properties are trivial (table 1). Periodic boundary conditions were applied to the RVE models as described earlier and the overall elastic properties of the stratum corneum were computed. This was done for two different cases – dry and wet stratum corneum. The final elastic properties of the stratum corneum for the two cases are shown in figures 11 and 12.

**Figure 11:**
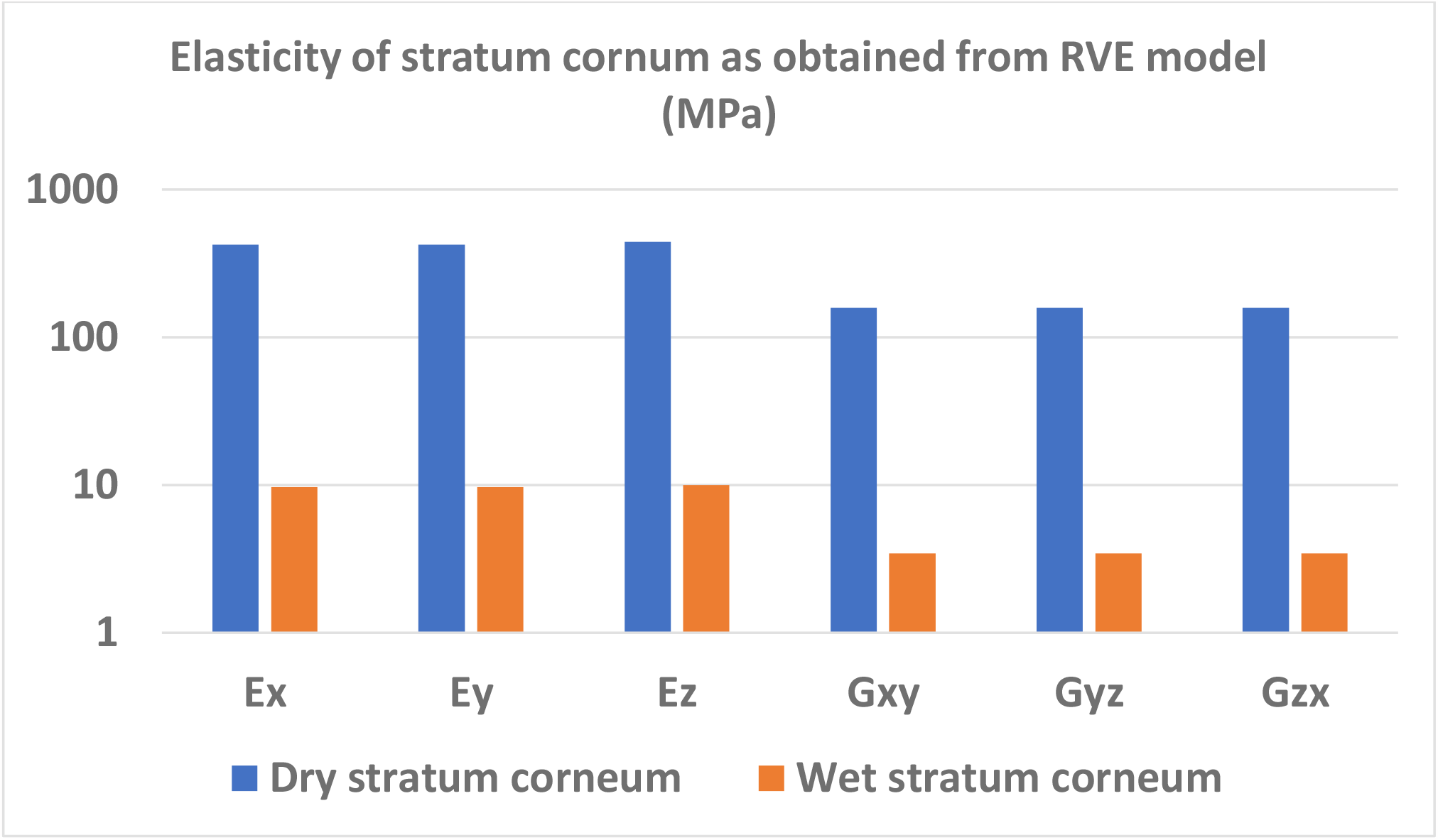
The elastic constants obtained as output from the RVE model for dry and wet stratum corneum.

**Figure 12:**
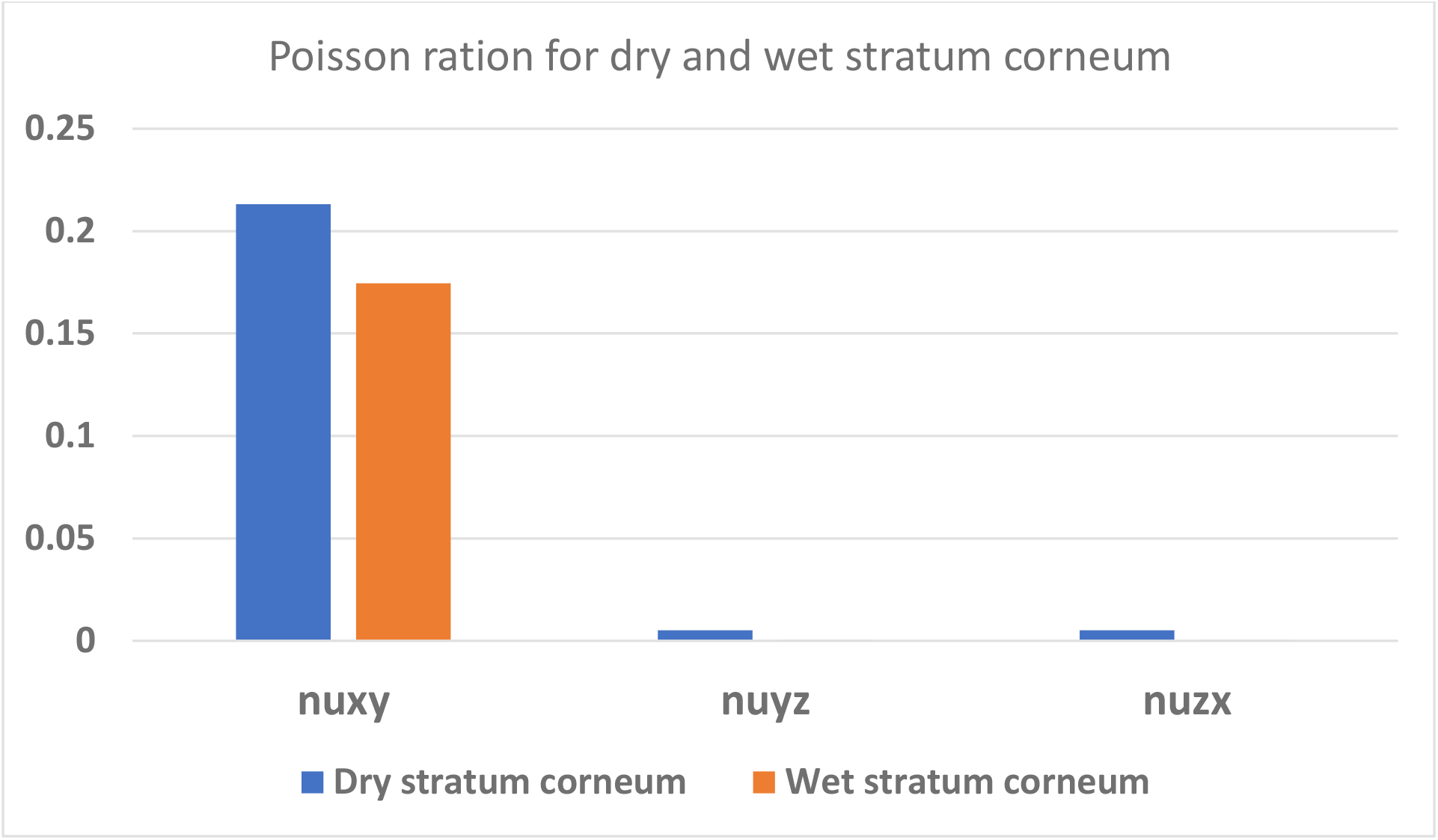
The Poisson’s ratio in the XY plane, which is the plane of the skin, for both dry and wet stratum corneum was found to be close to 0.25 which is in the same order of individual components’ Poisson’s ratio (0.2). However, Poisson’s ratio was very small for the other two planes as seen here. The reason for this might be the less thickness of stratum corneum RVE in Z direction. Since we are calculating the Poisson’s ratio computationally, the less thickness in Z direction might play a role while applying periodic boundary conditions to the RVEs leading to such a trend in results.

It is very commonly seen in the literature that increase in skin moisture decreases the skin stiffness and hence, reduces the elasticity parameters. Thus the RVE model can incorporate such variations in the skin mechanical properties. It should be noted that this drop in elasticity for wet stratum corneum was seen by only altering the properties in the intercellular space of stratum corneum. The properties of the corneocytes were left unaltered. This shows that despite the intercellular space occupying less than 10% of the volume in stratum corneum, it has a significant impact on defining the mechanical properties of the stratum corneum.

From figure 11, we note that the elastic parameters in each direction are almost similar, indicating isotropic behavior of stratum corneum. However, in figure 12, we seen that for both, dry and wet stratum corneum, the Poisson’s ratio in YZ and ZX plane is very small compared to XY plane. The reason for this might be the less thickness of stratum corneum RVE in Z direction. Since we are calculating the Poisson’s ratio computationally, the less thickness in Z direction might play a role while applying periodic boundary conditions to the RVEs leading to such a trend. However, the Poisson’s ratio in XY plane is close to 0.2 as expected since every component within the RVE has the same Poisson’s ratio.

The mechanical properties of stratum corneum obtained from the RVE model are then included in the multilayer skin model so that the effect of stratum corneum microstructure on the contact mechanics can be studied. The snapshot of the implementation is shown in figure 13. It can be seen that all the three models (two RVE and one multilayer skin) are connected to each other, and the generated data is transferred from first step to the last.

**Figure 13:**
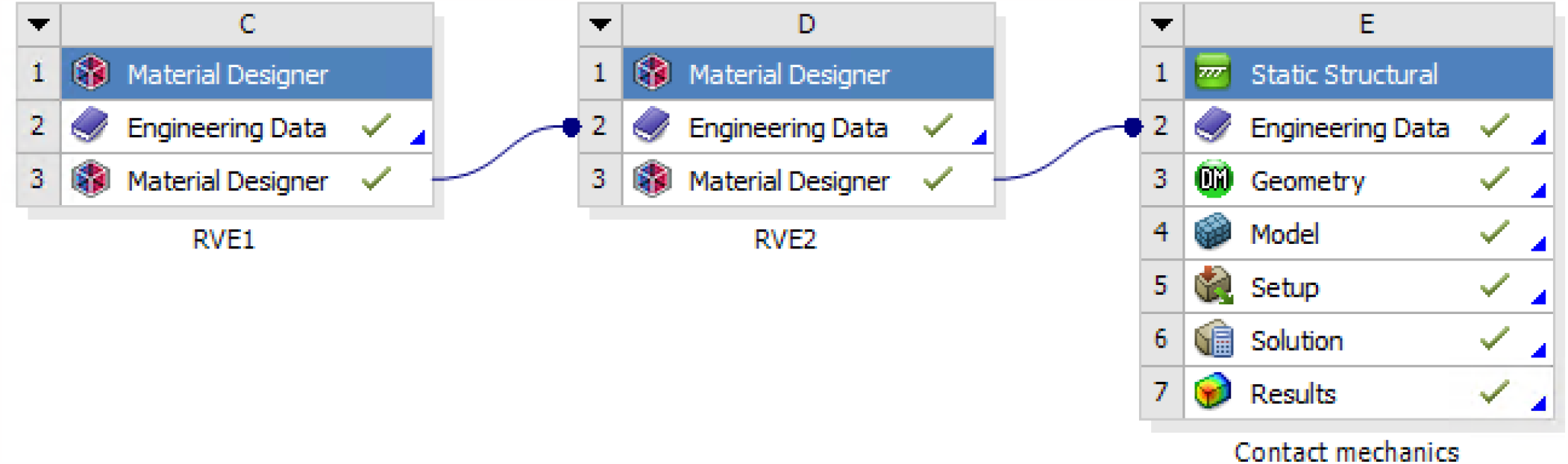
The complete framework connecting the RVE models with the contact mechanics (skin friction) model using Ansys 19.2. From the first RVE, the homogenized properties of a cellular scale layer are obtained. These layers are then stacked over each other to form a complete structure of the stratum corneum, and the homogenized properties of stratum corneum are computed in the second RVE. At the end, these properties of stratum corneum are fed into a multi-layer skin contact mechanics model.

### 3.2 Contact Mechanics Simulations

Using the RVE based finite element framework described in the previous sub-section, we simulate the indentation and sliding of a spherical indenter on a multilayered skin. The stresses and deformations predicted by the simulation are used to obtain the frictional stress between the indenter and the skin surface. By considering the properties of the microconstituents corresponding to the dry and wet stratum corneum conditions, the friction between skin and indenter is characterized for various conditions.

#### Geometry and meshing

Figure 14 shows the geometry of the simulation domain containing multilayer skin and indenter in Ansys 19.2. In the FE implementation, shown in figure 14, the thicknesses for various skin layers are as follows: stratum corneum: 0.02 mm, viable epidermis: 0.05 mm, dermis: 0.84 mm and hypodermis: 2 mm. The diameter of the indenter was kept 5mm. We used quadratic mesh elements. The meshing in stratum corneum was kept finer with around 30,000 elements. In the rest of the skin layers, there were around 900 elements. And finally for the sphere, there were close to 1500 elements. Each skin layer had 4 elements in the thickness direction.

**Figure 14:**
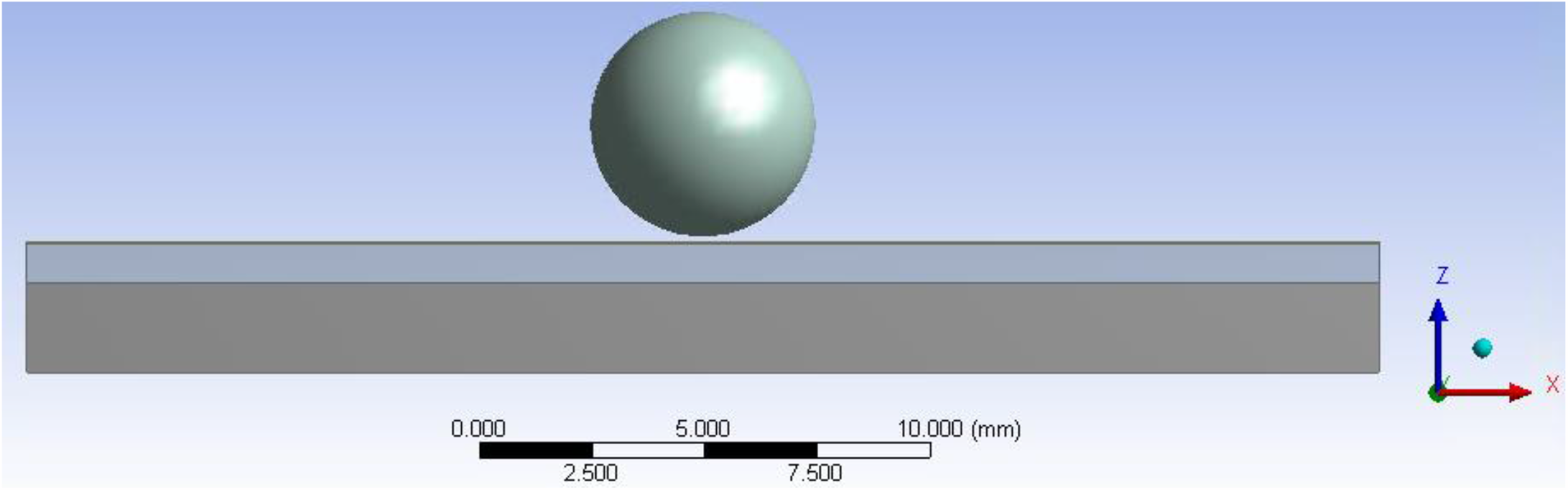
The contact simulation was done in Ansys 19.2, between a multi-layer skin model and a spherical indenter of fixed diameter. The skin sub-layers were ‘bonded’ together so that there was no slip between layers. The indenter was given displacement in two steps, first in -Z direction (indentation) and then in X direction (sliding) with different local COFs. Frictional force was computed as a function of indentation depth, stratum corneum elasticity, and local COFs.

#### Constitutive model parameters for other skin layers

We took the material properties of the stratum corneum from the previous RVE models and for the next three layers, we used the available data from literature (mentioned in table 2) (Diosa, Moreno, Chica, Villarraga, & Tepole, 2021).

**Table 2:** The mechanical properties of the three skin layers below the stratum corneum were taken from the literature. The stratum corneum’s material parameters were taken from the RVE model. Rest of the layers were considered Ogden first order hyperelastic materials with parameters (refer to equation 1) as mentioned in the table.

| Skin layer | Parameter 1 (MPa) | Parameter 2 |
| --- | --- | --- |
| Viable epidermis | $\mu = 7.74823$ | $\alpha = 2.4291$ |
| Dermis | $\mu = 0.0205$ | $\alpha = 3.2999$ |
| Hypodermis | $\mu = 0.0094$ | $\alpha = 4.0319$ |

#### Loading and Boundary conditions

The bottom-most face of the skin was fixed in position using ‘fixed support’ boundary condition. The skin layers were assumed to be bonded together in order to avoid slip between them. This was imposed using ‘bonded’ contact definition in Ansys. The indenter was given displacement in two steps: indentation into the skin up-to a certain depth and then a horizontal traverse. The horizontal motion was kept fixed at 2 mm. However, two cases of indentation depths were considered: 0.2 mm and 0.5 mm. For each case, the local COF between the stratum corneum and the indenter was also varied as discussed previously. Using the contact tool in Ansys, the frictional stress colormap plots were obtained for each case.

### 3.3 Frictional stress colormap plots

The contact mechanics simulations were run by considering various indentation depths and local coefficients of friction (COF) for stratum corneum properties corresponding to dry and wet skin conditions. The following sections discuss about the contact frictional stress colormap plots at the end of horizontal traverse step for different cases.

#### Comparison of frictional stresses for dry vs wet stratum corneum

**Figure 15:**
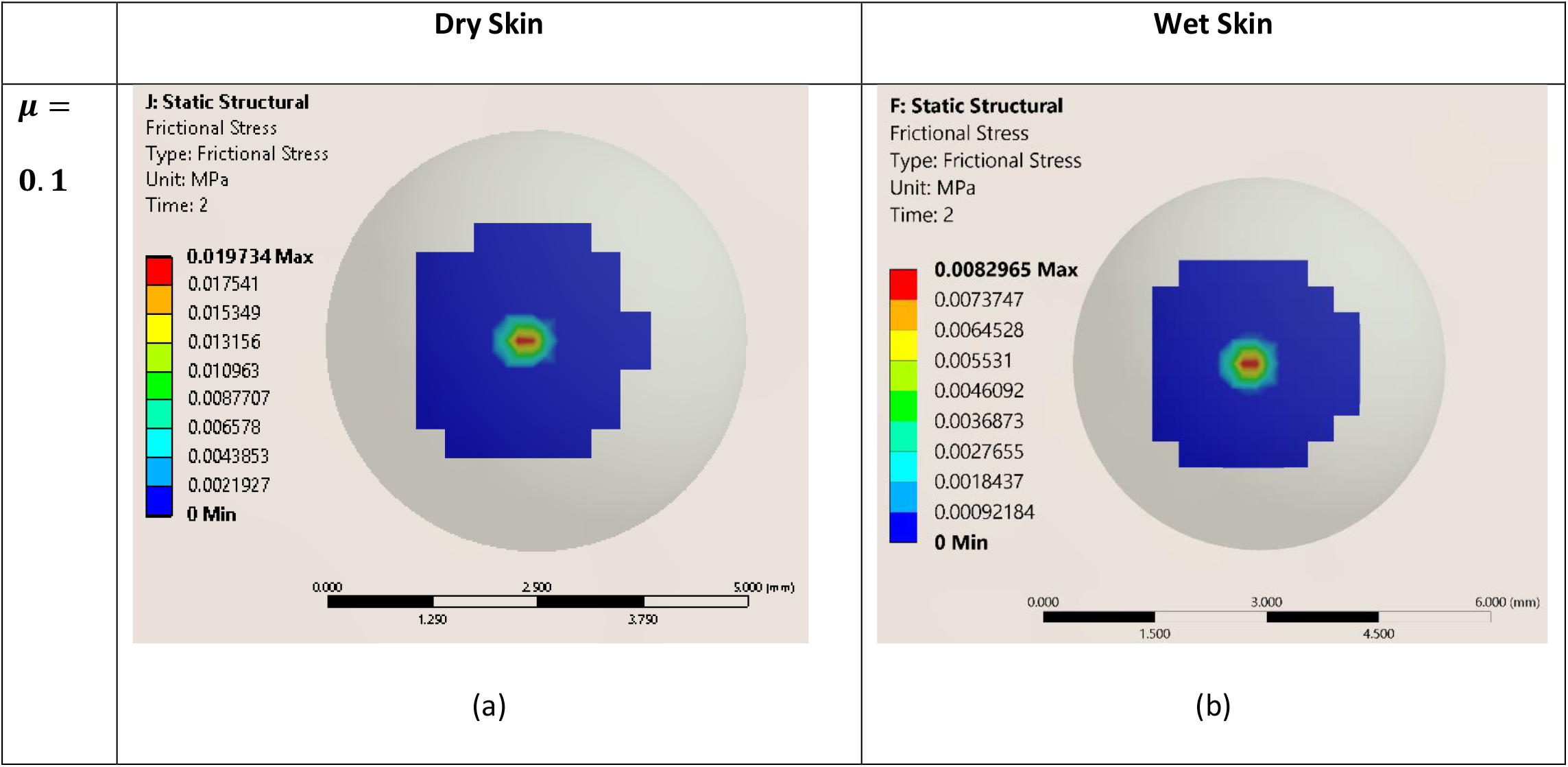
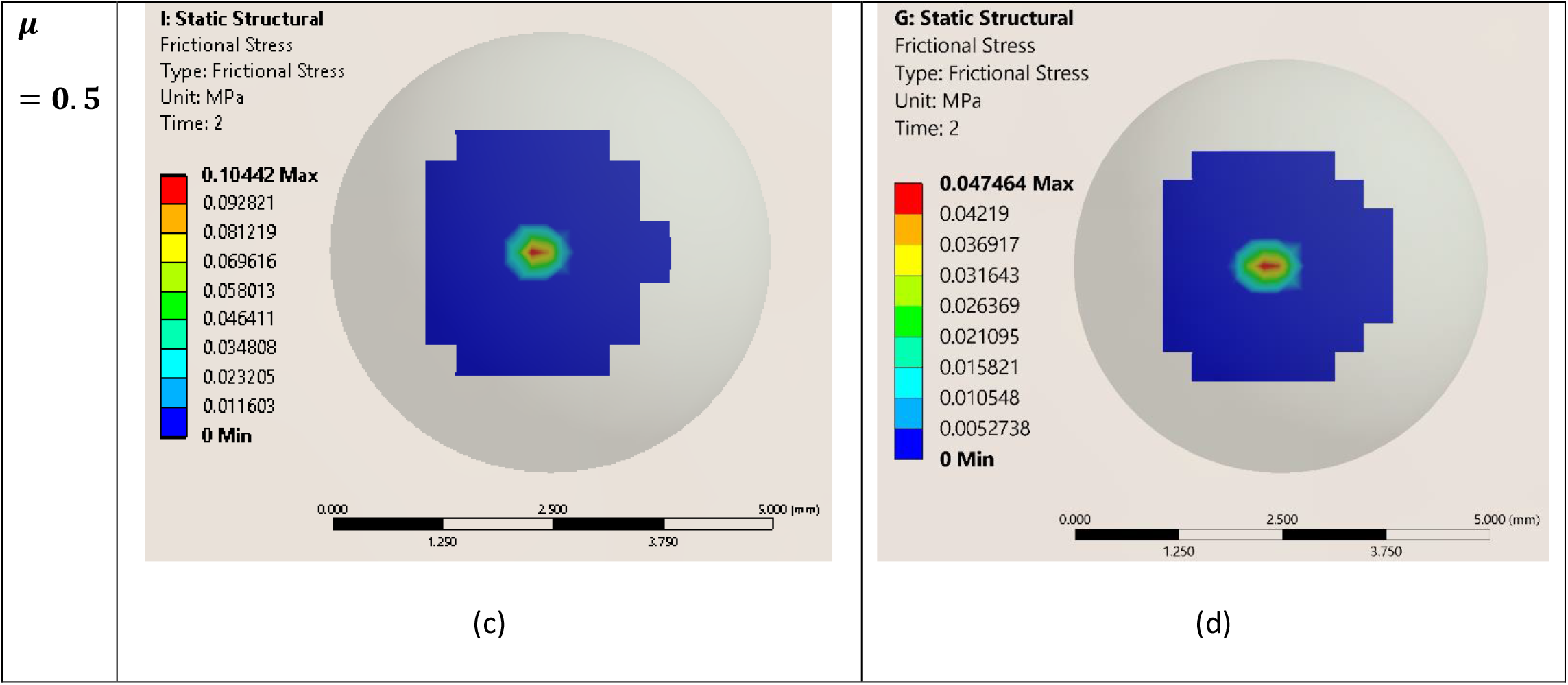
Frictional stress colormap plots of dry and wet stratum corneum at 0.2 mm indentation depth and different local coefficients of friction (μ): a) Dry Stratum Corneum, μ= 0.1, b) Wet Stratum Corneum, μ= 0.1, c) Dry Stratum Corneum, μ= 0.5, Wet Stratum Corneum, μ= 0.5.

Using the current framework, the effect of moisture on the stratum corneum can be implemented by changing the elasticities and volume fractions of its micro-constituents in the RVE model. We observe that dry stratum corneum is considerably stiffer than the wet stratum corneum and the intercellular space between the corneocytes play a major role in determining the two cases. A higher stiffness in dry stratum corneum results in more resistance to motion between the skin and the indenter. Hence, the frictional stress observed for dry stratum corneum is higher than wet stratum corneum. In previous experimental works, the authors have compared the frictional force observed in dry and wet skin and found that increasing the humidity in skin initially increases the frictional force, but it starts to decrease after a point (Derler, Rossi, & Rotaru, 2015). This increase in frictional force has been attributed to molecular interactions between water and skin and surface energies (Kim & Yun, 2019). For high humidity in skin, these interactions lead to increase in contact area (van Kuilenburg, Masen, & van der Heide, 2015) and adhesive forces (Pailler-Mattei & Zahouani, Analysis of adhesive behaviour of human skin in vivo by an indentation test, 2006) between the skin and the indenter, which increases the frictional force (Gerhardt, Strässle, Lenz, Spencer, & Derler, 2008; Kim & Yun, 2019). These kinds of interactions were not incorporated in the current model and so the frictional stresses solely depend on the stratum corneum’s elasticity. It should be noted that even though the thickness of stratum corneum (0.02 mm) is considerably smaller than the overall skin thickness (3 mm), it still plays a major role in contact mechanics.

The RVE models can be further improved by understanding the impact of water on the stratum corneum at molecular scale. Similar RVE models can also be obtained for lower layers of skin such as dermis and hypodermis which can give an even better control over defining properties of skin. The physical attributes of the skin also vary with age, gender, anatomical location etc., and these changes can be accommodated for in the two RVE models or in the multilayer skin model.

#### Effect of indentation depth on the frictional stress

**Figure 16:**
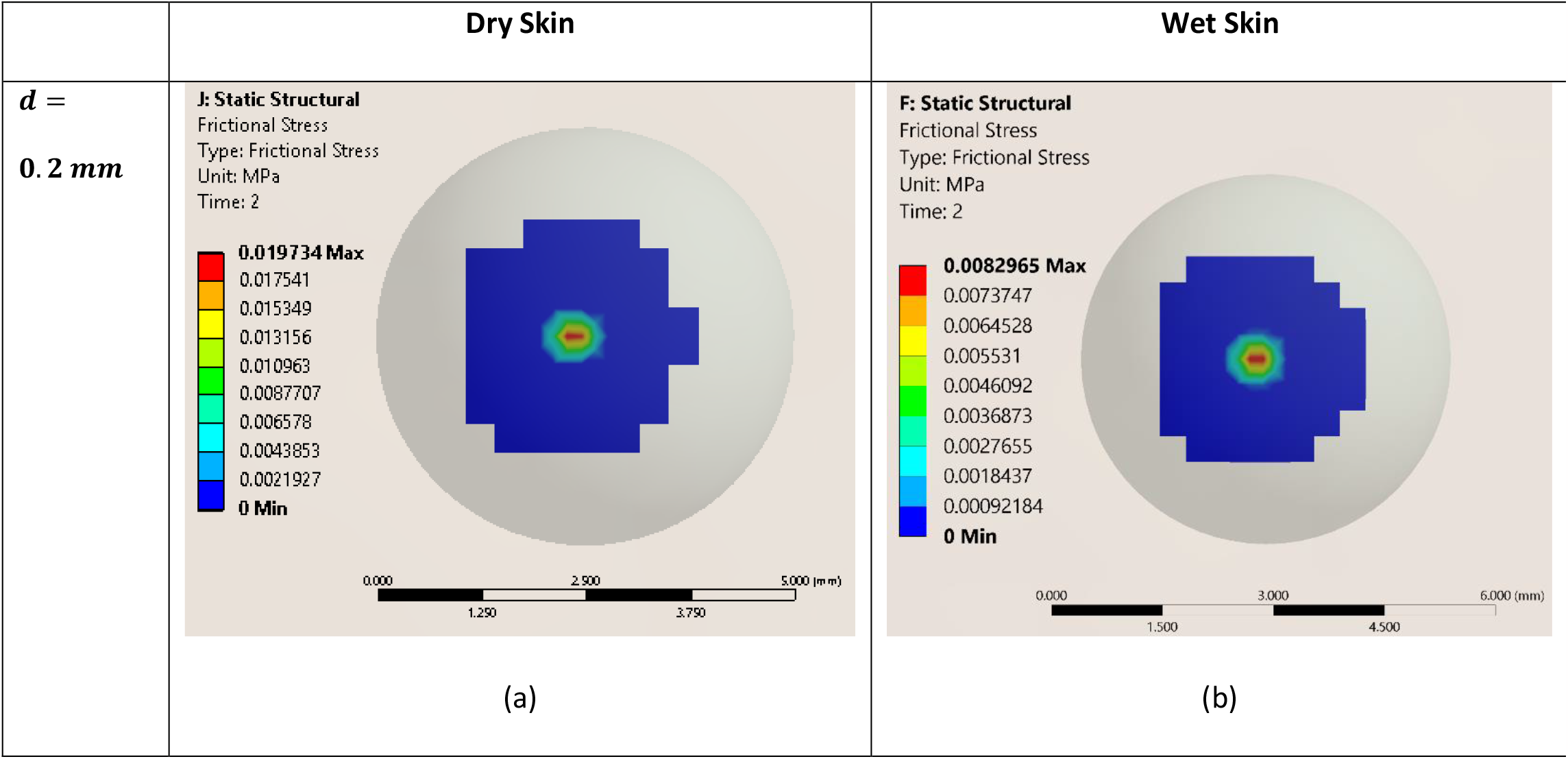
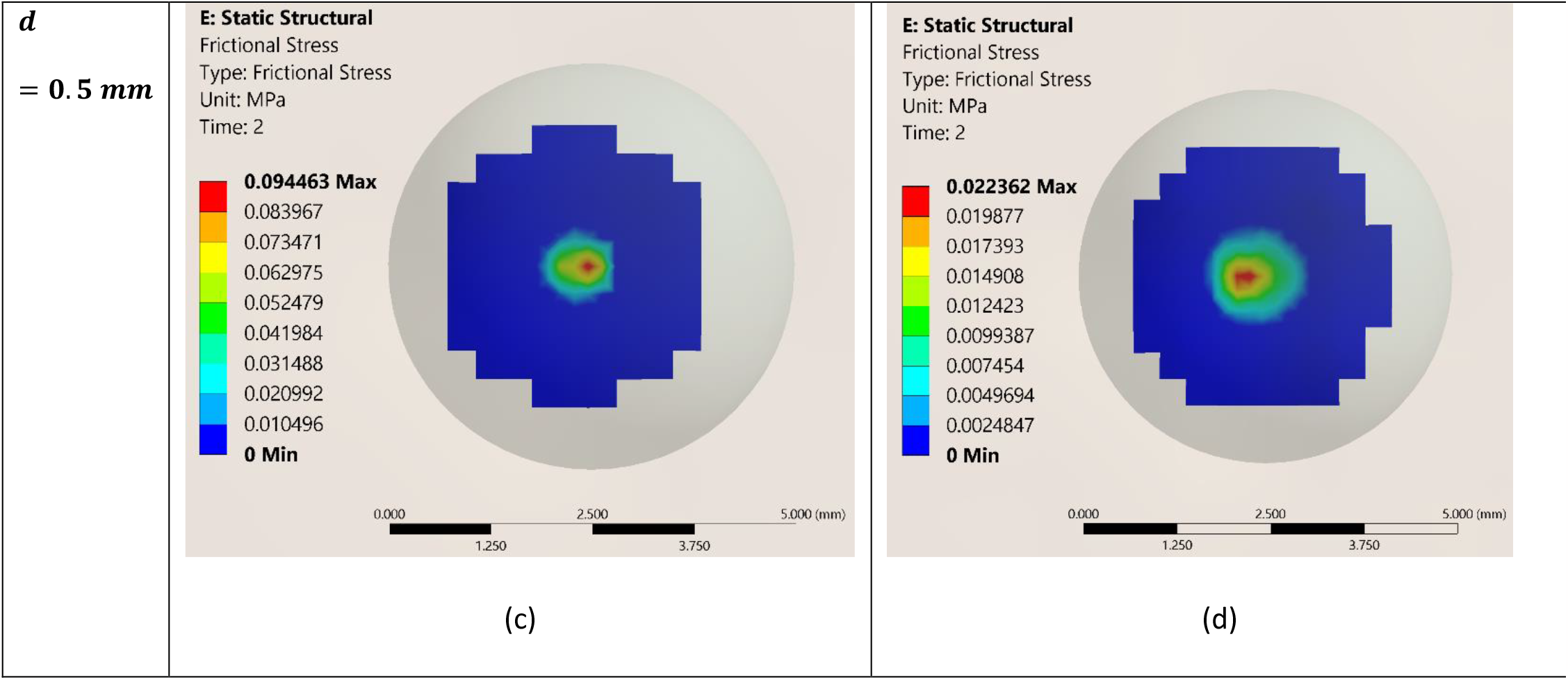
Frictional stress colormap plots of dry and wet stratum corneum at local coefficient of friction of 0.1 and various indentation depths (d): a) Dry Stratum Corneum, d = 0.2 mm, b) Wet Stratum Corneum, d = 0.2 mm, c) Dry Stratum Corneum, d= 0.5mm, d) Wet Stratum Corneum, d= 0.5mm.

We ran simulations for two different indentation depths – 0.2 mm and 0.5 mm. Naturally, a lower indentation depth will result in a lower contact area and vice versa. For higher indentation, since there is more skin surface resisting the motion of the indenter, we observe a higher frictional stress. This can be attributed to the deformation component (Greenwood & Tabor, 1958) of friction in soft material. Friction in soft materials also depends on the contact area. Since a higher indentation means a higher contact area, the increase in frictional stress is further justified. Previous works on in-vivo skin indentation tests have shown that the frictional force increases with increase in normal load of indentation (Adams, Briscoe, & Johnson, 2007; Pailler-Mattei & Zahouani, Analysis of adhesive behaviour of human skin in vivo by an indentation test, 2006), which in turn is directly proportional to the indentation depth (Pailler-Mattei & Zahouani, Analysis of adhesive behaviour of human skin in vivo by an indentation test, 2006). These results indicate that the current work can be used to study the impact of indentation depth on the skin’s contact mechanics with any foreign object.

#### Effect of local coefficient of friction on the frictional stress for dry skin

**Figure 17:**
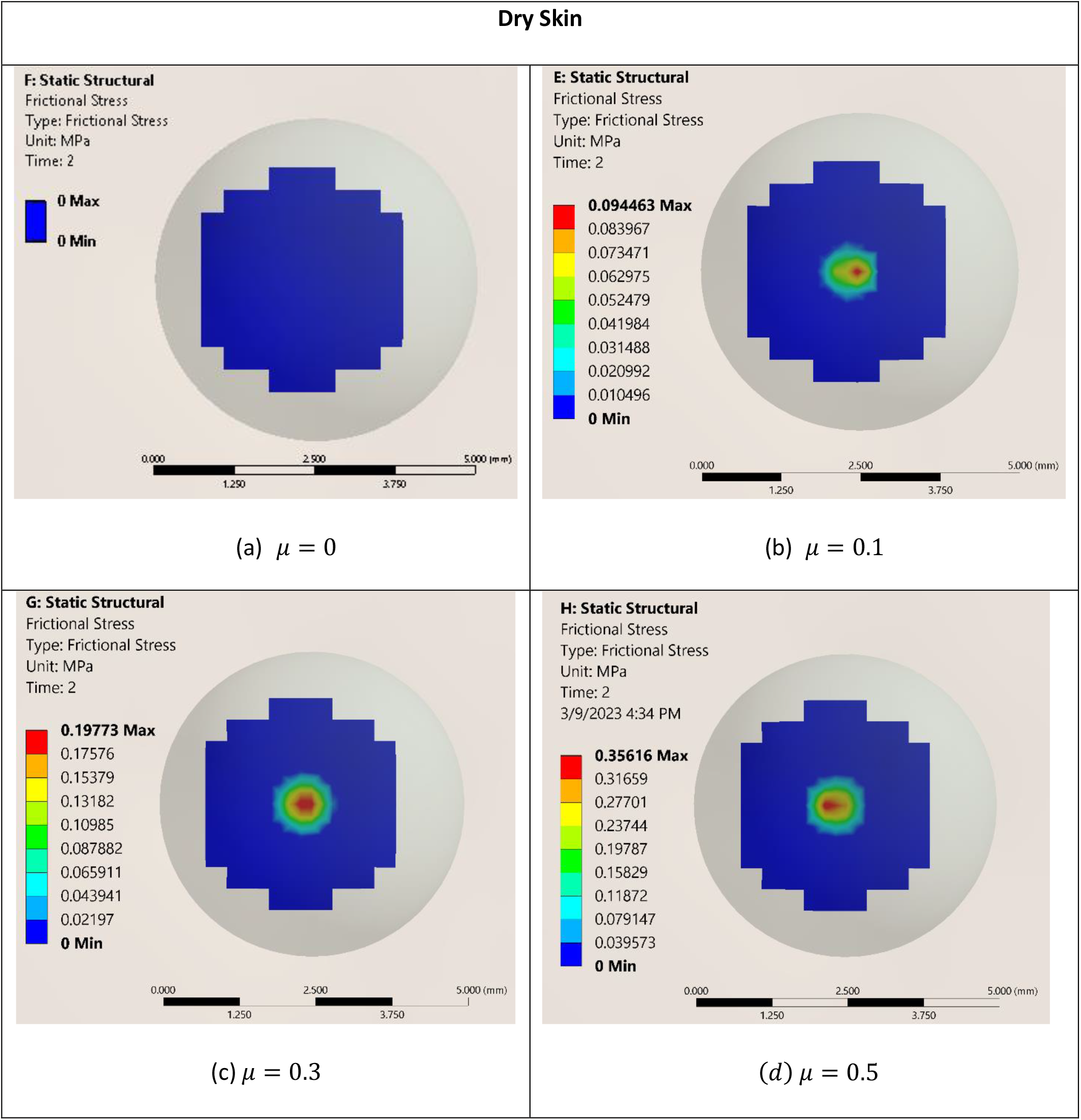
The frictional stress colormap plots for dry stratum corneum (E = 350 MPa) at 0.5 mm indentation depth for different local coefficient of frictions ((a) = 0, (b) = 0.1, (c) = 0.3, (d) = 0.5)

The local COF between the skin and any indenter depends on surface properties of both (Adams, Briscoe, & Johnson, 2007; Pailler-Mattei & Zahouani, Analysis of adhesive behaviour of human skin in vivo by an indentation test, 2006; Dzidek, Bochereau, Johnson, Hayward, & Adams, 2017). Usually, for a very smooth indenter like steel, the local COF would be small. However, for a rough material like rubber, the local COF might be more. In order to accommodate for this difference in properties, we varied the local COF between 0 and 0.5. It was observed that there is a slight increase in maximum frictional stress when the local COF is increased. This increase indicates more resistance to motion of the indenter or more frictional force. It has been observed that indeed for a combination of rough indenter (rubber) and skin, the frictional force is higher compared to smooth indenter (steel) and skin (Adams, Briscoe, & Johnson, 2007; Dzidek, Bochereau, Johnson, Hayward, & Adams, 2017). Hence, this model can be used for variety of such indenter-skin combinations by changing the local COF. The local COF definition can be further improved by implementing the topology of the skin surface (roughness) and adhesive forces between the two contacting surfaces.

In the contact mechanics model, we have considered all the layers of skin to be perfectly flat. This works fine if the indenter size is much larger compared to the micro-asperities on skin surface. However, we can also impart a surface roughness to the top layer of skin to get a better picture of skin’s frictional properties as implemented in previous works (Leyva-Mendivil, Lengiewicz, Page, Bressloff, & Limbert, 2017). Also, we have used a spherical indenter in our study. We can easily use any other geometry of the indenter in the contact mechanics framework to study the friction as a function of indenter geometry.

The idea behind this framework was to reduce the use of experiments to determine the frictional properties. Skin friction depends on a number of factors (age, race, moisture etc.) and it would require a lot of experimental simulations and data gathering, in order to develop empirical relations. This framework can provide an alternative to generate that amount of data by simply varying the modeling parameters pertaining to the microstructural components of skin. That data can be then used by a learning algorithm which can further aid in the design of devices, fabrics, or any such object and optimize for functionality and comfort level.

### 3.4 Contact Pressure Comparison

The nature and degree of contact between the skin and some indenter can be quantified by measuring the contact pressure at the area of contact. We did a comparison study of the contact pressure observed at the end of the indentation step for dry and wet stratum corneum for different indentation depths (0.2 mm and 0.5 mm).

#### Contact pressure colormap plots for dry and wet stratum corneum

**Figure 18:**
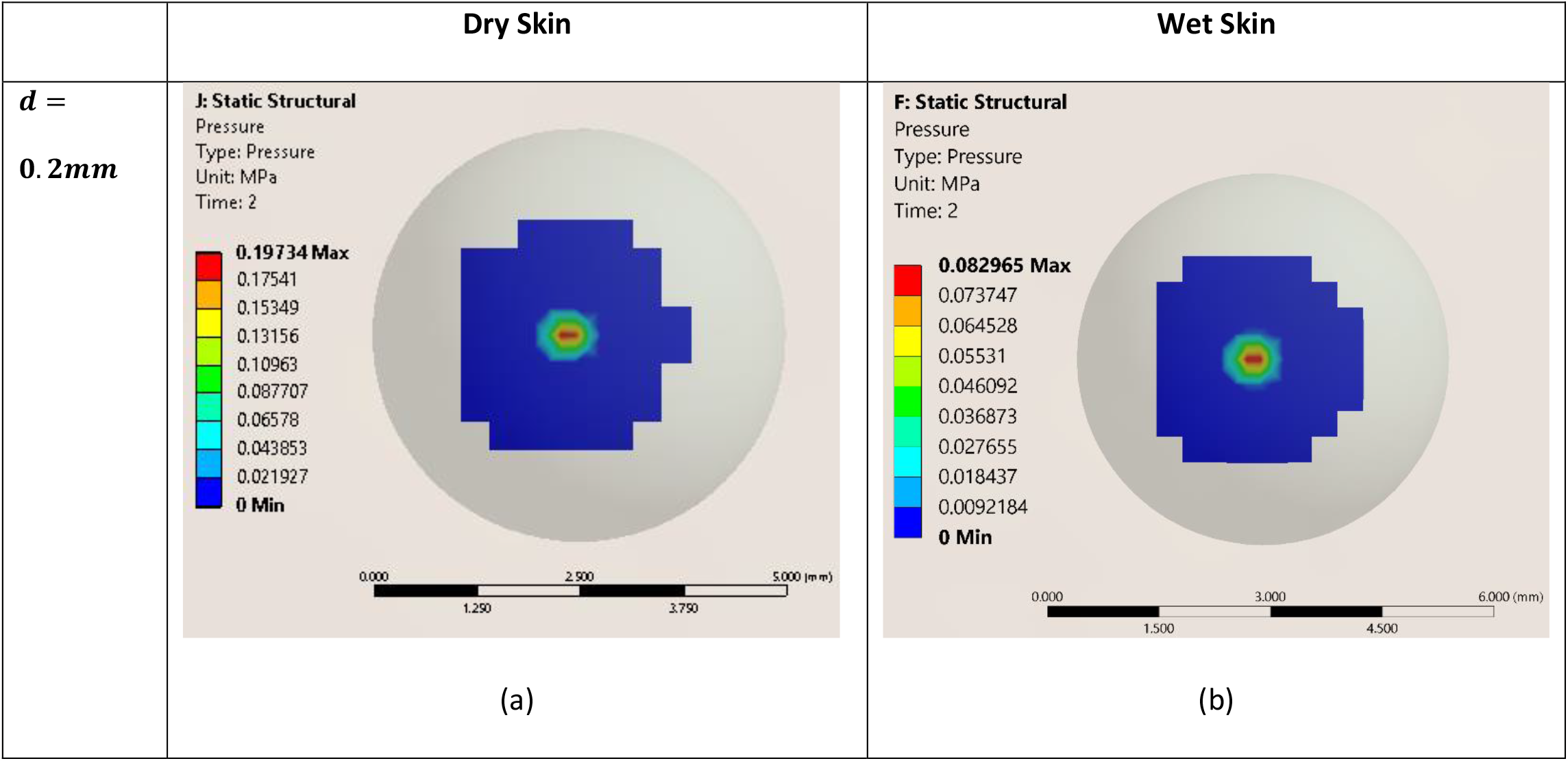
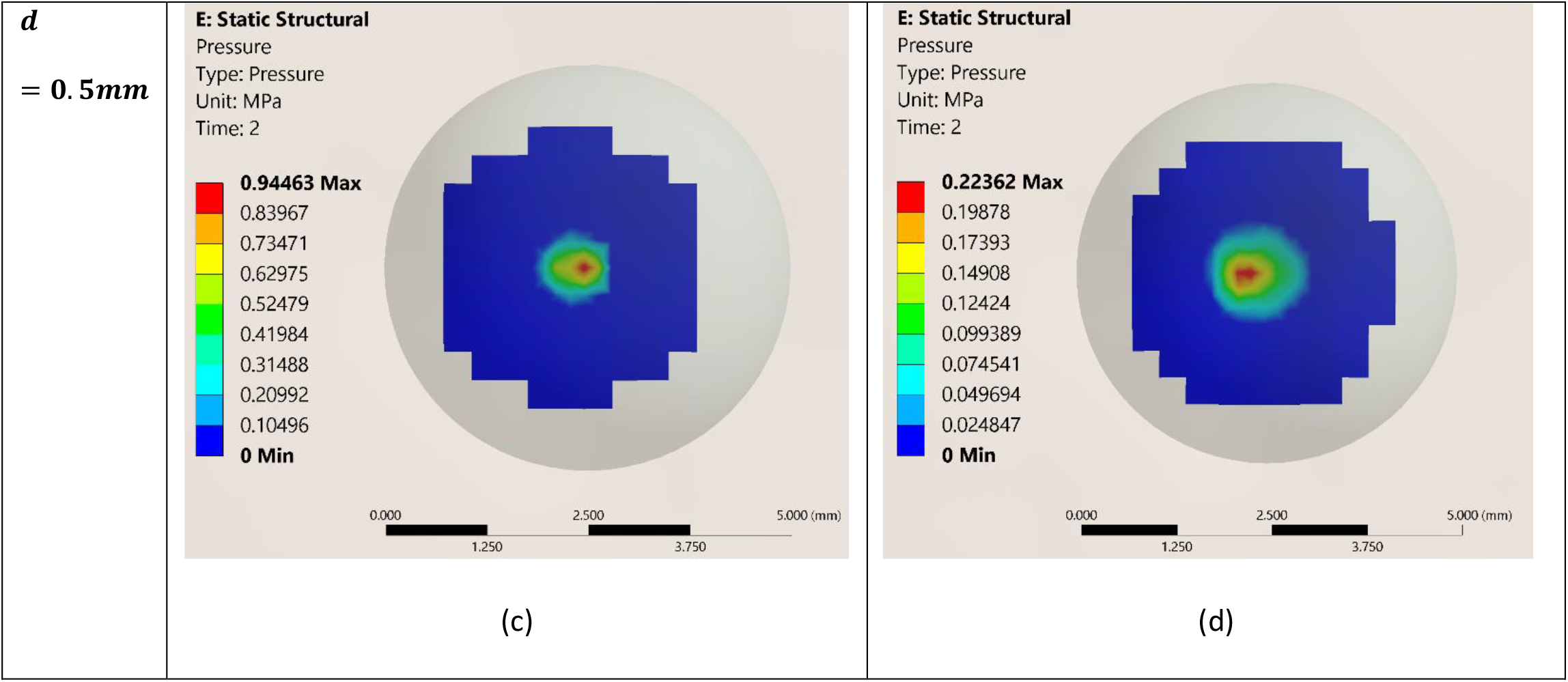
Contact pressure colormaps of dry and wet stratum corneum at local coefficient of friction of 0.1 and various indentation depths (d): a) Dry Stratum Corneum, d = 0.2 mm, b) Wet Stratum Corneum, d = 0.2 mm, c) Dry Stratum Corneum, d= 0.5mm, d) Wet Stratum Corneum, d= 0.5mm.

Since the dry stratum corneum is much stiffer than wet stratum corneum, we observe that the maximum contact pressure for dry stratum corneum is higher. The contact pressure for higher indentation is distributed over a larger area indicating an increase in contact area. According to the Hertzian theory of contact in soft materials (Johnson, Kendall, & Roberts, 1971), the maximum contact pressure increases with increase in indentation depth. However, we only observed a slight increase in the contact pressure for higher indentation. This may be due to the anisotropic, stratified structure of the skin, whereas the Hertzian theory holds for linear isotropic materials only. Also, the Hertzian theory is usually used for micro-indentation tests, whereas the dimensions of the two contacting bodies in our case are in the same order of magnitude.

## 4. Conclusions

In this work, we have simulated contact mechanics between skin and indenter using an RVE model for stratum corneum, which plays the most significant role in contact mechanics for small indentations. The RVE model shows that the intercellular space, although occupying a significantly less volume, plays a major role in determining the stratum corneum’s mechanical properties. Dryness or wetness of the stratum corneum can be simulated in-silico by changing the microstructural properties of the stratum corneum, using the RVE model. The contact mechanics model takes in the stratum corneum properties as input from the RVE model and we see that, despite a very low thickness, the stratum corneum contributes a lot in determining frictional properties. Dry stratum corneum being stiffer than the wet stratum corneum provides more resistance to the motion of the indenter and hence more frictional stress. We also see that with increase in indentation depth or local COF, the maximum frictional stress increases. Increasing depth increases the deformation component of friction whereas the local COF captures the surface properties of the indenter, a smoother surface like steel has less local COF than a rough surface like rubber. Hence, the current framework provides an in-silico alternative, to study contact between skin and any foreign object and variety of data can be generated by simply changing the control parameters like skin geometry/thickness, indenter geometry, skin microstructure and so on. The data generated from this finite element modelling framework can be used to train ML models that can be used to conduct systematic parametric studies with lesser computational power. This approach has been explored by researchers in the recent past (Yin, Zhang, Yu, & Karniadakis, 2022) and will be part of our future work.

## Acknowledgements

The authors would like to thank Susmit Sahu for his support in running the simulations and post-processing. The authors would also like to thank K Ananth Krishnan, CTO, Tata Consultancy Services for his constant encouragement and support during this project.

